# Divining Deamidation and Isomerization in Therapeutic Proteins: Effect of Neighboring Residue

**DOI:** 10.1101/2021.07.26.453885

**Authors:** Flaviyan Jerome Irudayanathan, Jonathan Zarzar, Jasper Lin, Saeed Izadi

## Abstract

Deamidation of asparagine (ASN) and isomerization of aspartic acid (ASP) residues are among the most commonly observed spontaneous post-translational modifications (PTMs) in proteins. Understanding and predicting a protein sequence’s propensity for such PTMs can help expedite protein therapeutic discovery and development. In this study, we utilized proton-affinity calculations with semi-empirical quantum mechanics (QM) and µs long equilibrium molecular dynamics (MD) simulations to investigate mechanistic roles of structure and chemical environment in dictating spontaneous degradation of asparagine and aspartic acid residues in 131 clinical-stage therapeutic antibodies. Backbone secondary structure, side-chain rotamer conformation and solvent accessibility were found as three key molecular indicators of ASP isomerization and ASN deamidation. Comparative analysis of backbone dihedral angles along with N-H proton affinity calculations provides a mechanistic explanation for the strong influence of the identity of the n+1 residue on the rate of ASP/ASN degradation. With these findings, we propose a minimalistic physics-based classification model that can be leveraged to predict deamidation and isomerization propensity of therapeutic proteins.

## Introduction

Protein stability is at the center of biological mechanisms spanning molecular pathways and therapeutic interventions.[1-3] Proteins are subject to various post-translational modifications (PTM) that are essential for their function, e.g. glycosylation and acylation. Some of these PTMs are spontaneous chemical reactions that lead to degradation, e.g. deamidation of asparagine (ASN→ASP/iso-ASP), isomerization of aspartic acid (ASP→iso-ASP) and oxidation of MET/TRP residues.[4, 5] These spontaneous chemical modifications often serve as a key indicator of stress that elicits a physiological response.[6] Bio-manufactured therapeutic proteins such as peptides, vaccines, monoclonal and bispecific antibodies (mAbs), and viral capsids are subject to physical and chemical stressors during manufacturing that result in accumulation of various PTMs.[7-11] Spontaneous chemical modification can negatively impact the biomolecule’s therapeutic efficacy, quality and developability by altering the structure and/or molecular properties of the protein. For instance, isomerization of an ASP residue alters the backbone connectivity and deamidation of an ASN residue imparts a negative charge (Figure1), both of which can result in altered functional consequences.[12, 13] Often protein engineering strategies are employed to mitigate the negative impacts of such chemical modifications, but this process is often iterative which prolongs the development cycle of a successful therapeutic.[14-16] Understanding and predicting the propensity of a therapeutic protein to undergo chemical degradation under stress can be leveraged to expedite development timelines.[17-20]

**Figure 1.**
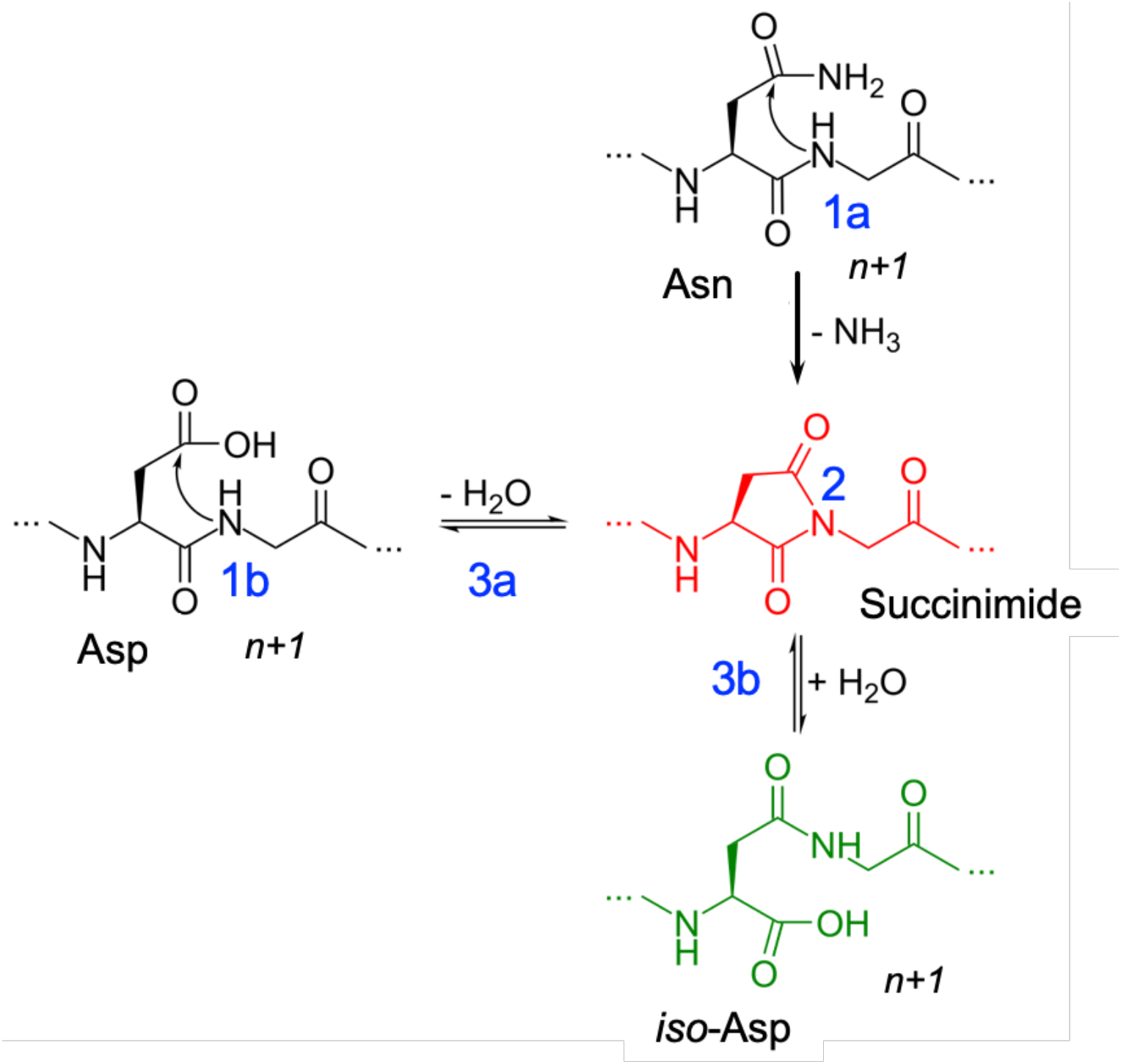
Deamidation (1a) and Isomerization (1b) reaction mechanisms showing deprotonation (1) of the n+1 amide leading to the formation of succinimide intermediate (2) and hydrolysis of succinimide (3) into aspartic acid (3a) or iso-aspartic acid (3b).

Deamidation of an ASN residue has been experimentally well characterized.[21-23] Among the multiple pathways that are feasible, pH-dependent base-catalyzed nucleophilic attack has been shown to be the predominant pathway.[23-26] The reaction is irreversible and proceeds via (Figure 1): 1) Base-catalyzed deprotonation of the n+1 amide 2) nucleophilic attack of the anionic nitrogen on the side chain carbonyl group and a subsequent ring closure that leads to the formation of a succinimide intermediate and 3) hydrolysis of the succinimide into ASP or iso-ASP. Similarly, the isomerization reaction can proceed from an ASN or ASP residue via the same succinimide intermediate pathway. There are mechanistic parallels and kinetic dissimilarities between the isomerization and deamidation reactions.[13, 27, 28] A comprehensive investigation of both reactions has revealed that the side chain of the neighboring n+1 residue can explain fold differences in the 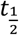 of the reaction in peptides.[21, 29-32] Previous studies using pentapeptides by Robinson et al.[33] have shown a higher likelihood of chemical degradation if the n+1 neighboring residue of the ASN residue is a GLY, ALA, SER or THR. Based on this, the sequences NX and DX (where X could be G/A/S/T) are often considered as sequence hotspots in proteins. However, these peptide hotspots do not sufficiently predict the reaction propensity in 3D folded proteins; a recent experimental survey of 131 clinical stage therapeutic (CST) antibodies (henceforth referenced as the Adimab dataset) by Lu et al. [34] revealed vast discrepancies in the range of deamidation and isomerization propensities among antibodies that do not follow the hotspot rules. For instance, of the twenty-seven NG hotspots only fourteen underwent chemical degradation. Similarly, of the forty-four DG cases only sixteen were isomerized. In addition, chemical degradation was observed at the non-hotspot residue locations such as NY, NW, DR and DY. The discrepancy reported in Lu et al. indicates a gap in our knowledge of how neighboring residues contribute to the chemical degradation propensity.

Multiple computational strategies, from physics-based approaches (e.g. quantum mechanics/molecular mechanics (QM/MM) and Molecular Dynamics (MD) simulations) [29, 30, 35, 36] to machine-learning models,[17, 20, 37] have been developed over the past two decades to predict deamidation and isomerization in peptides and proteins. Several detailed computational models explored site specific effects using QM/MM calculations at the peptide level. For example, ab initio calculations on model compounds have revealed the contribution of backbone secondary structure affecting the proton-affinity (Figure 1a) of the n+1 amide.[38] However, it is as of yet unclear if these insights are generalizable to ASP and ASN residues with varied neighboring residues on a large set of proteins. Another noteworthy study[36] used MD and QM/MM to compute conformational and chemical free energy barriers along the deamidation reaction pathway. While this study provided detailed insights into the free energy changes, the method was only tested on a small number of degradation sites on a single molecule and thus would be computationally prohibitive for high throughput screening. Recently, multiple machine learning models have been proposed that were trained with a number of structure-based features such as secondary structure, local flexibility, size of the neighboring residue, and solvent exposure.[17, 20, 37, 38] While computationally facile and practical in nature, these models often ignore important mechanistic and structural details (such as conformational dynamics) that are important for the reaction’s feasibility. Besides, it remains unclear how broadly generalizable these models are, given that they were trained on a limited set of data points.

Here we have utilized microsecond long equilibrium molecular dynamics (MD) simulations and proton-affinity calculations to understand the impact of neighboring residues in dictating the chemical degradation at ASP/ASN site of proteins. Based on the analysis of a large set of 131 CST antibodies, we propose a physics-based classifier model that can accurately predict the propensity of deamidation and isomerization of a protein sequence. The model relies on decomposing the major steps of the reaction (deprotonation, nucleophilic attack and hydrolysis) as conformational degrees of freedom and inferring the reaction propensity based on the thermodynamic feasibility of exploring those conformations.

## Results and Discussion

### Conformational Determinants of Deprotonation

#### QM calculations and crystal structure analysis

Deprotonation of the amide nitrogen has been experimentally characterized to proceed via a base-catalyzed reaction that is quite spontaneous even at acidic conditions.[31, 39-41] Several studies have investigated the dependence of backbone amide acidity on the conformation and chemical environment using QM/QMMM calculations.[29, 30, 35-37] In one of the pioneering works, Radkiewicz et al. [26, 27] used N-formyl-glycinamide as a model compound representing a peptide bond to calculate the relative proton affinity as a function of the ϕ and ψ dihedral angles (using HF/6-31+G*//HF/3-21G). This analysis showed that the acidity of backbone amide is largely impacted by of the ψ dihedral angle; ϕ angle was found to have a small effect. Here, we carried out DFT calculation to estimate proton affinity for the same model compound at the M062X/6-311++G(d,p) level of theory in both gas phase and aqueous phase continuum SMD (Solvation model based on density) solvent environment. Our results are in agreement with the earlier studies: proton affinity of backbone amide deviates significantly with respect to changes in the backbone conformation, Figure 2b and the SI. Specifically, when ϕ and ψ are at angles observed within α-helices, the proton bonded to the amide experiences a repulsive force from the adjacent carbonyl carbon as a result of a di-electric instability with the amide nitrogen (Figure 2c) i.e. proton affinity of the model compound is lower in the α-helical conformation centered around -60°≤ψ≤60°, Figure 2b. The gas phase semi-empirical QM calculations using MOPAC with PM6-D3H4,47 PM7 and RM1 Hamiltonians (see methods and the SI) are also consistent with DFT results.

**Figure 2.**
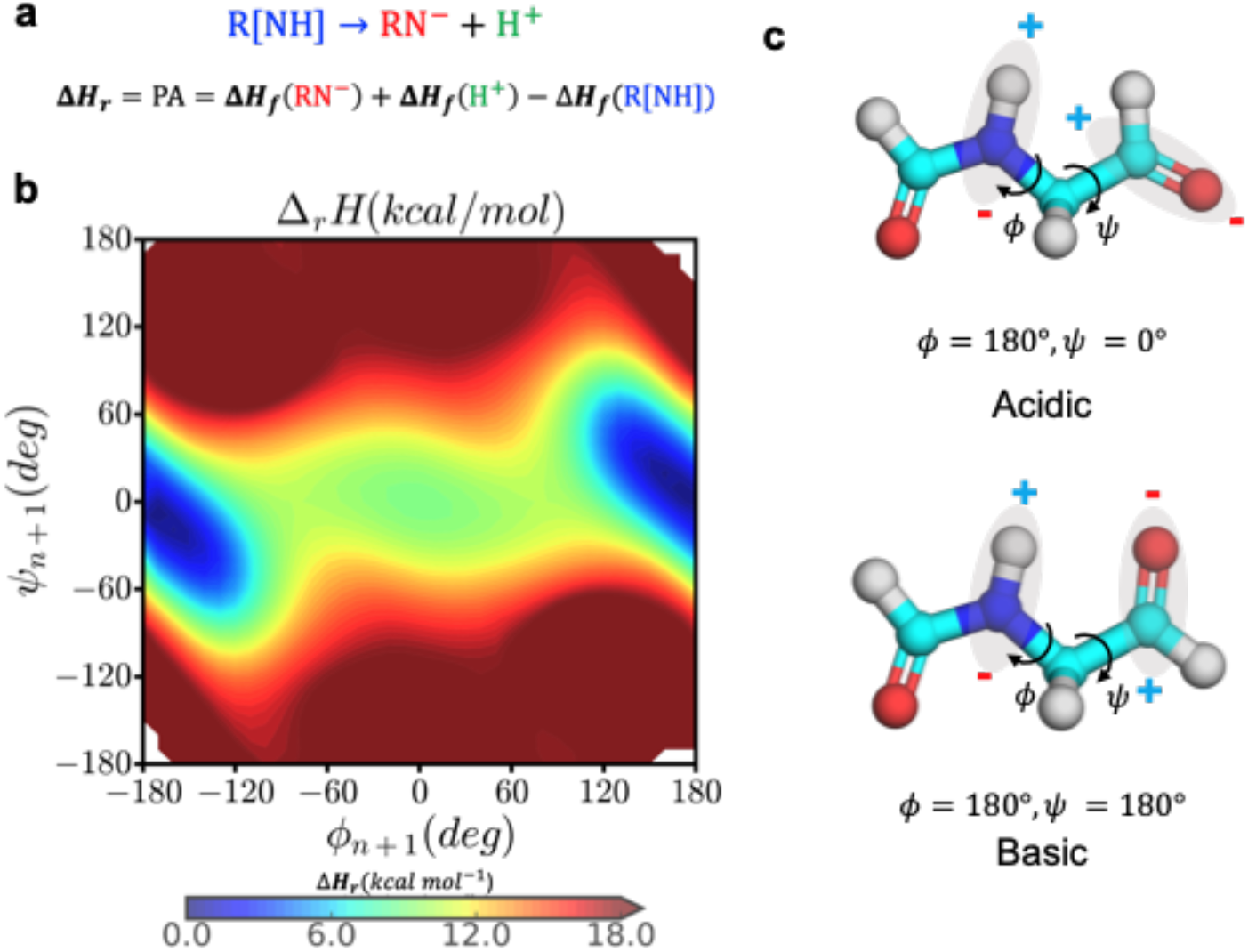
Backbone conformation dependence for proton affinity. (a) the deprotonation reaction and the calculation of proton affinity as the heat of formation of the forward reaction, (b) the proton affinity distribution of the small molecule N-formyl-glycinamide as a function of the backbone dihedral angles, and (c) the most acidic and basic conformations based on the proton affinity distribution.

We then sought to find the generalizability of this conformation-dependent amide acidity rules in the context of peptides. In order to do so we extended the proton affinity calculations to DX and NX dipeptides as a function of their backbone ϕ & ψ dihedral angles, Figure 3 and the SI. The goal was to investigate if proton affinity calculations can explain the fold differences of reaction half-lives observed in pentapeptides due to the variations in the n+1 residue. For instance, an interesting comparison can be made between Gly and Ala as n+1 residue. In solution, GGNGG and GGNAG peptides show high propensity to adopt an α-helical conformation[41, 42] yet their deamidation half-lives are an order of magnitude different (1.03 and 21.1 days respectively).[33] Our dipeptide proton affinity analysis suggests that the amides in both NG and NA are acidic dependent on the ψ angle, -60°≤ψ≤60° being acidic (Figure 3a). However, contrary to the model compound data, the proton affinity of the amide in the NA peptide significantly changes as a function of the ϕ angle. Specifically, the backbone amide of the NA residue is more acidic in conformations when ϕ>0 compared to when ϕ<0 within the same boundary of -60°≤ψ≤60°. This indicates that the proton affinity NA peptide is also dependent on the type of the helical conformation (right-handed α-helix vs left-handed α-helix). A right-handed α-helix conformation places the methyl sidechain group in close proximity to the backbone amide that offers protection from exposure to the base and/or hydrolysis (Figure 3c, conformation 2). Similarly it can be inferred from the proton affinity plots (see SI) that: 1) the sidechain mediated protection increases with increasing the size of non-polar, hydrophobic groups (i.e. Phe > Ile > Leu > Val > Ala), and 2) the presence of a polar hydroxyl or carbonyl groups in the n+1 side-chain (as in Asp, Ser, Thr, Asn, Tyr) can facilitate the deprotonation of the amide by solvent mediated electrostatic interactions. In the case of GLY or in any left-handed α-helix conformation the protection from the n+1 residue is completely lost (Figure 3c, conformation 3).

**Figure 3.**
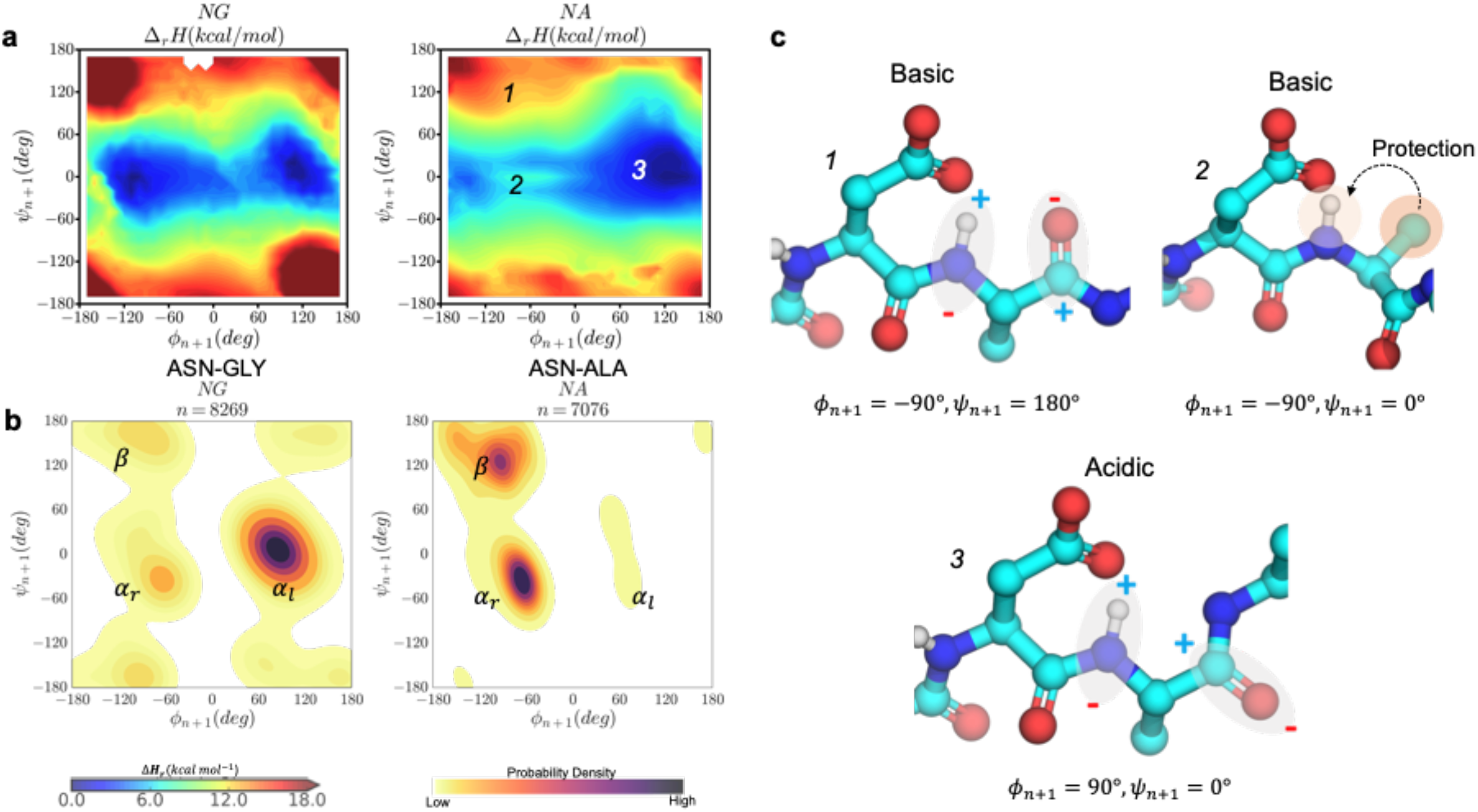
Conformational determinants of proton affinity in NG and NA dipeptides. a) the gas-phase proton-affinity calculated with the semi-empirical RM1 Hamiltonian as a function of ϕ & ψ angles. The colors scale from 0→18+ kcal/mol from blue to red. b) the corresponding kernel density estimation plot of the respective dipeptides observed in the PDB database of crystallized antibodies. The color gradient scales from low→high from yellow to purple. c) representative basic and acidic conformations beta sheet, right-handed helix and left-handed helix.

It must be noted that a large fraction of combinations of ϕ & ψ dihedral angles explored in the dipeptide models are energetically inaccessible in folded proteins. To identify the relevant secondary structures for the NX and DX motifs in proteins, we collected ϕ & ψ angle distribution from all the antibody crystal structures available from the PDB database. These distributions were analyzed using kernel density estimation (KDE) to arrive at the probability density function (PDF) of conformational preference for [N/D]G and [N/D]A residues in ϕ & ψ space (Figure 3b and SI). Comparing the proton affinity calculations with the conformational PDF reveals that [N/D]G sites mainly populate the acidic left-handed α-helical conformations, whereas [N/D]A sites largely favor the basic right-handed helix (ϕ<0 region) and a beta sheet. The latter is in agreement with a previous work showing that a steric interaction between the methyl group and the terminal carbonyl oxygen destabilizes all structures with ϕ∼120°, relative to glycine dipeptides.[43] These results clearly explain why [N/D]A sites are often orders of magnitude less prone to degradation than [N/D]G sites.

A comprehensive analysis of the proton affinity calculations and conformational PDF in ϕ & ψ space for all the [N/D]X sites (SI) revealed that the frequency of observing a residue in the acidic region from the PDB is strongly correlated with the likelihood of the residues to undergo degradation. In particular, we observed that the most frequently modified [N/D]X sequences based on various publications (i.e. DG,[44-46] DS, [14, 47, 48] DD,[48] DT,[20] DH,[49] NG, NS, NN, and NT[12, 20, 33, 44]) mainly favor the right-handed helix in the Ramachandran plot, i.e. the more acidic region. Interestingly, the strongest right-handed helix population is seen for DG and DS, followed by DD, DN and DT. Similarly, for deamidation motifs, a decreasing population of right-handed helix is observed in the order of NG, NS, NN and NT. These results correlate with the approximate hierarchy of these hotspots based on published studies.[34]

Overall, these observations indicate that the ϕ & ψ backbone dihedral angle of the peptide bond at an [N/D]’s n+1 position can be used as a metric to identify sites that have higher propensity to deprotonate: 1) β-sheet conformation is less deprotonation prone, 2) the left-handed α-helix regions (−60°≤ψ≤60°, ϕ>0°) is acidic for all amino acids that have a non-polar (A, V, L, I) or bulkier side-chain groups (F, W), 3) On the contrary for small polar amino acids (G, S, T, D) both the left and right-handed α-helix (−60°≤ψ≤60°) conformations are acidic.

#### Generalizability of the conformation dependent backbone acidity in proteins

To test the generalizability of the proton affinity data at the peptide-level, we explored the relationship between backbone conformations of all of the [N/D]X sites in the CDR region of 131 therapeutic monoclonal antibodies and their reported degradation rates. To obtain the backbone conformation preference, we calculated the conformational free energy surface (FES) in the ϕ & ψ space for the n+1 residue from microsecond long equilibrium molecular dynamics (MD) simulations of the Fab structures. We arbitrarily chose a value of 2KbT energy barrier or ∼1 kcal/mol to define energetically favorable conformations, i.e. in the FES a region whose energy values are < 1 kcal/mol are likely to be sampled under equilibrium conditions. This dataset corresponds to 1039 [N/D]X sites (498 ASP and 541 ASN) of which 30 DX and 39 NX sites were prone to degradation (reactive).

Figure 4 shows that the calculated conformational distribution (PDF) of reactive and non-reactive residues correspond very well with the expected conformational preference seen in proton affinity calculations. The experimentally reported non-reactive residues show preference to the right-handed α-helix or the β-sheet conformation while the reactive residues show large preference to the left-handed helical conformation. Figure 6 (D1) reveals a correlation between the rate of degradation and the probability of the backbone amide being found in the acidic regions. With an exception of two sites, the degradation rates fall below the desirable 5% cutoff for all the [N/D]X sites that are in a basic backbone conformation (i.e. Free Energy > 1kcal, which corresponds to either a beta-sheet or a right-handed helix when the n+1 residue is non-polar). However, the results in Figure 4 suggest that when the backbone is in an acidic conformation (Free Energy < 1kcal), the [D/N]X site is not necessarily guaranteed to undergo degradation. In other words, a high propensity for a conformation with a basic backbone hydrogen can prevent the site from degradation but lack thereof is not sufficient for degradation.

**Figure 4.**
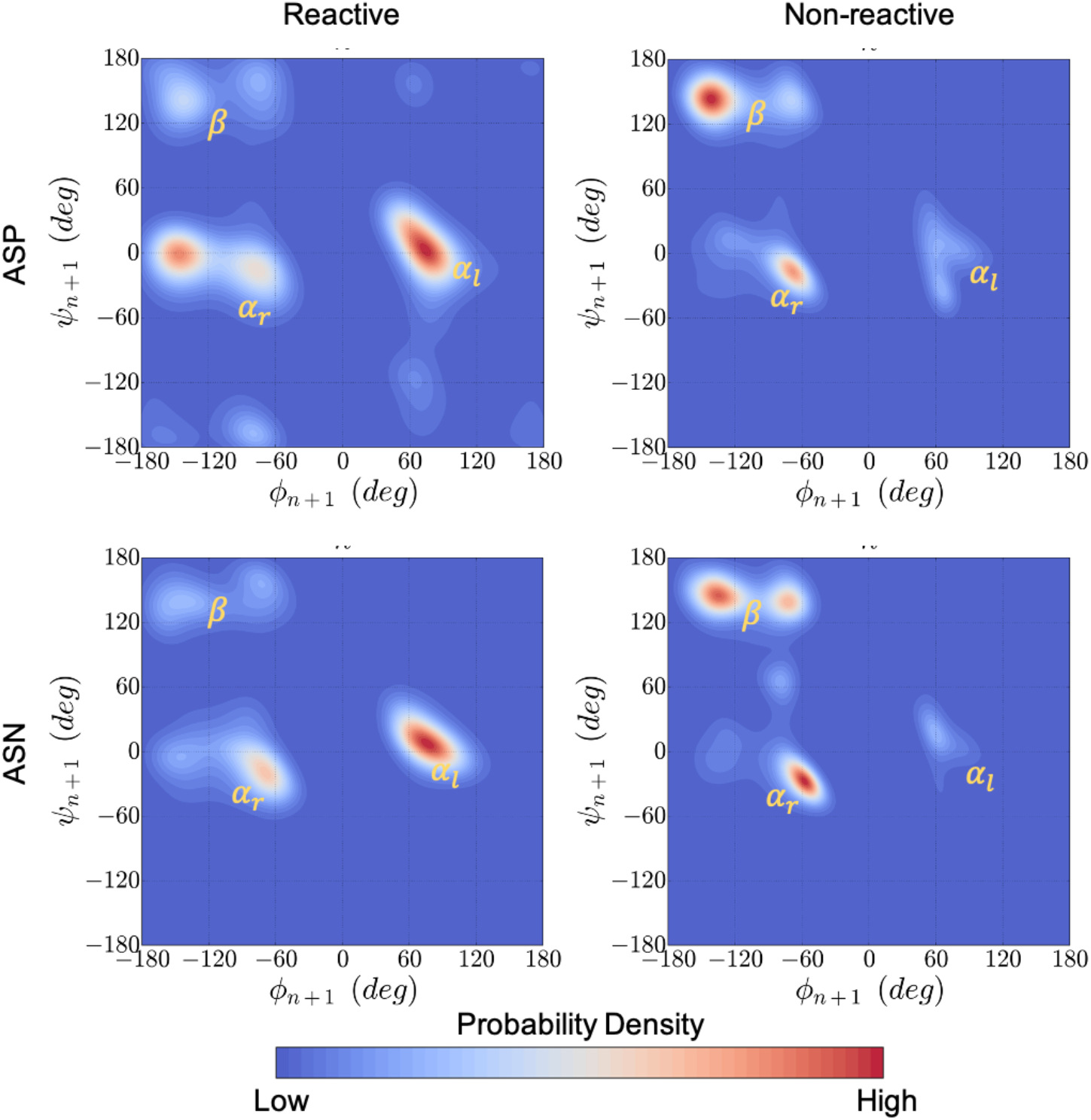
Kernel density estimation plots for the backbone conformation of all 1039 n+1 residues that neighbor an ASP (top) or ASN (bottom) in the adimab dataset. The ϕ & ψ angle distribution was collected from μs long equilibrium MD simulations and classified as reactive (left) and non-reactive (right) based on the report from the adimab set. A clear pattern emerges around the left-handed α-helix conformation in reactive cases that is distinct from the non-reactive cases on the right. The right most panel shows the Ramachandran plot boundaries labelled with the α-helix and β-sheet conformations.

Overall, the backbone conformational preference of NX and DX residues under equilibrium conditions as a single descriptor was found to classify the reactive vs non-reactive residues at 65% accuracy (Table 1). This shows that proton affinity inferred from the secondary structure of n+1 residue is a good descriptor to predict degradation propensity of NX and DX sites.

**Table 1.**
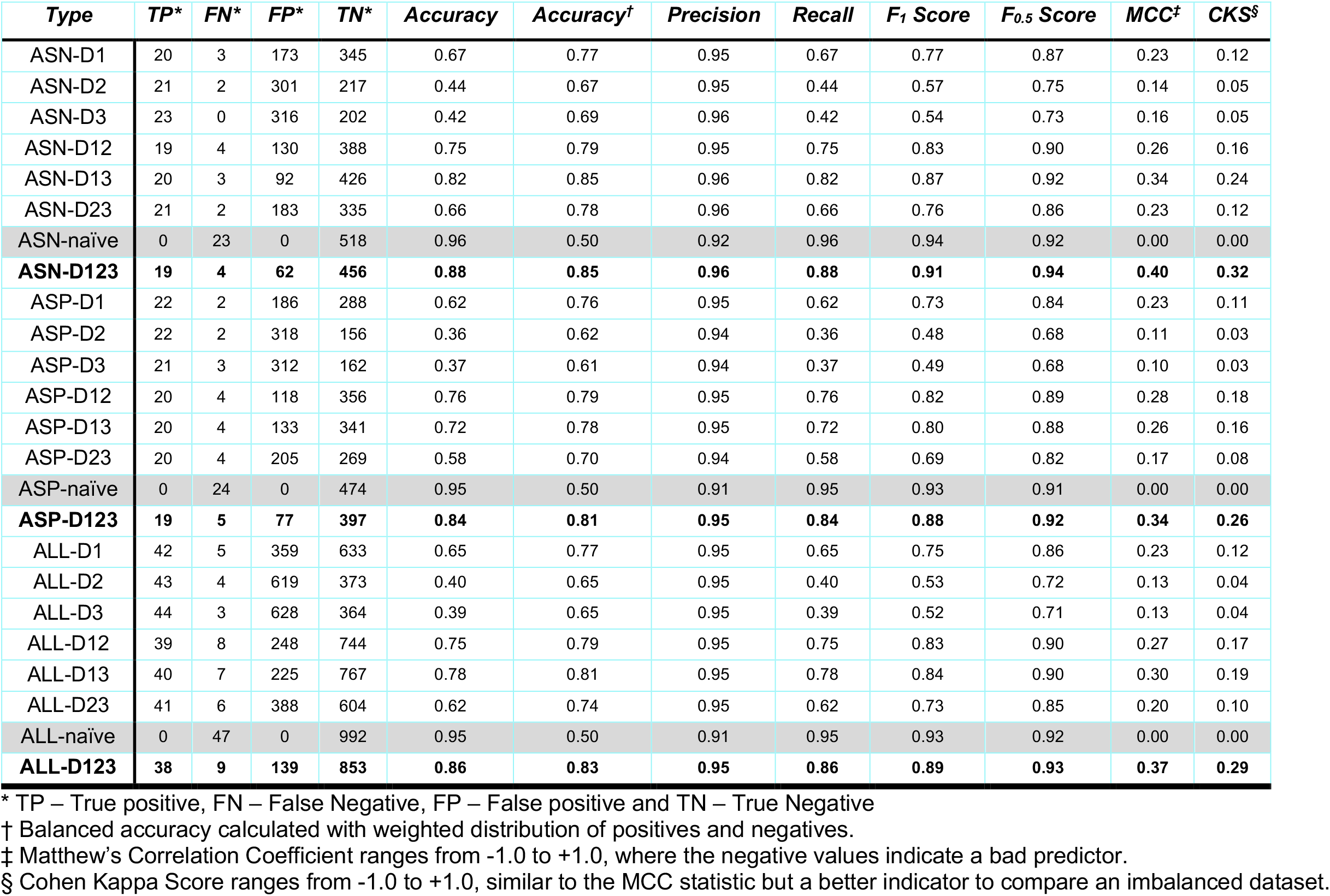
Performance statistic for the physics-based classifier model when different descriptors are used: D1-backbone dihedral conformation of the n+1 residue, D2-side-chain dihedral conformation of ASP/ASN residue, D3-fraction of time the ASN/ASP residue remains solvent accessible. Combinations are represented by DXX where the X represents the descriptors. The true distribution of the adimab set is 47 TP and 992 TN.

#### Conformational determinants of cyclization

Once deprotonation is enabled by the acidity of the backbone, the next step in the reaction is ring closure leading to succinimide formation via nucleophilic attack between N-atom of the n+1 residue and Cγ of the ASN/ASP residue (Figure 1). In order for the nucleophilic attack to be feasible the N- and Cγ should come closer than distances of 3.0 Å (near-attack conformation).[29, 30] In folded proteins sidechain interaction including hydrogen bond formation can limit the conformational flexibility of ASN/ASP residues to visit near attack conformations. This in turn can disfavor succinimide formation. Multiple sidechain orientations can result in the near attack conformation (Figure 5a). Conformational degrees of freedom affecting the distance between Cγ and N_n+1_ can be defined as a function of the two dihedral angles ψ & χ_1_.

**Figure 5.**
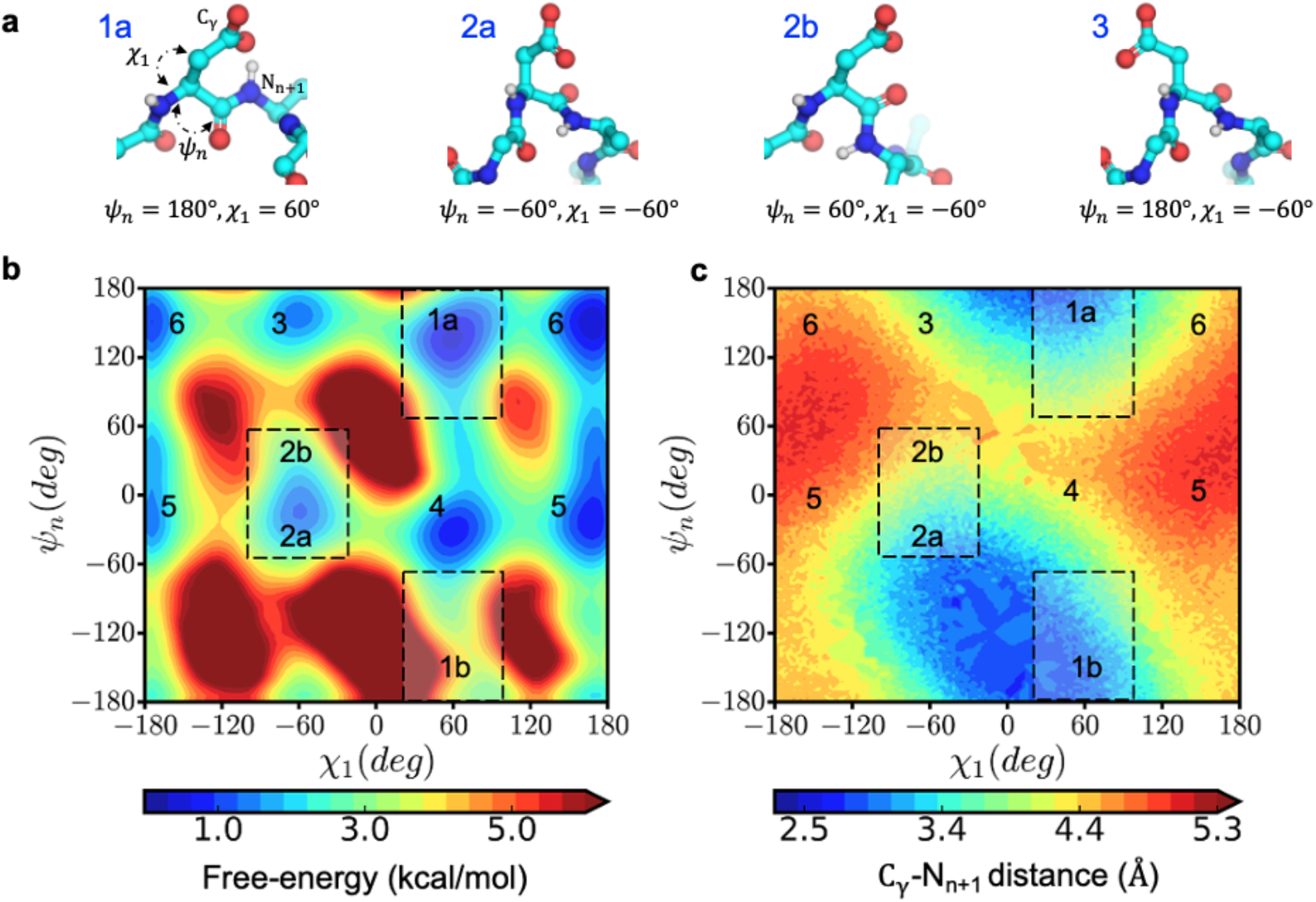
Conformational space for the near-attack conformation that enables succinimide formation. a) The structural orientation of the reactive (1a and 2a) and non-reactive sidechain conformations (2b and 3) of an aspartic acid residue is shown in ball and stick representation. b) Free-energy surface of the side-chain conformation in ψ & χ dihedral angle space and c) the corresponding distance between Cγ and the n+1 amide nitrogen calculated from metadynamics simulation of Ace-GGNAG-Nme pentapeptide.

**Figure 6.**
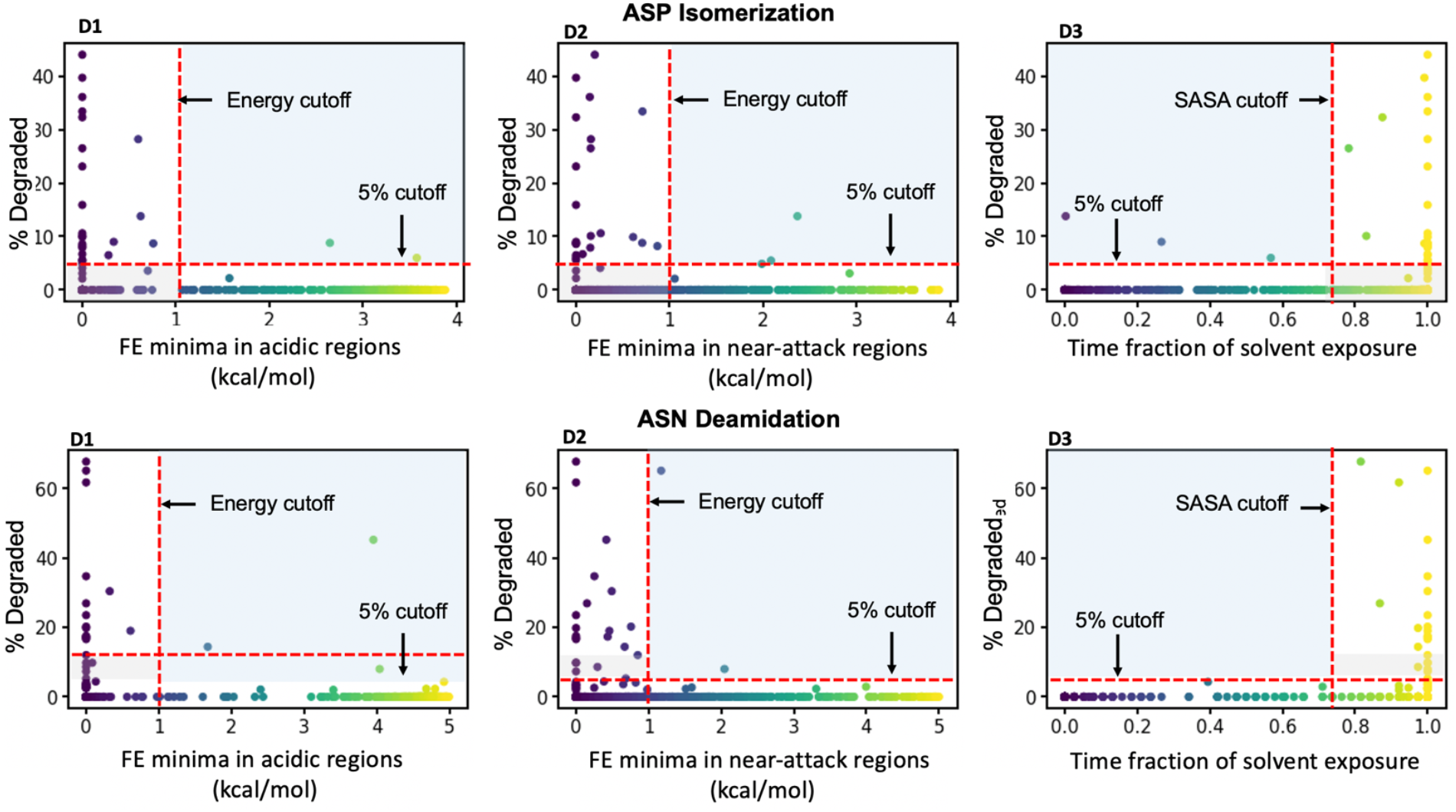
Performance of individual descriptors in discriminating degraded ASP (top) and ASN (bottom) residues from the adimab dataset. The percentage degradation values reported in the adimab study are represented on the X-axis. The Y-axis in the leftmost panel (D1) is the free-energy barrier at the deprotonation prone backbone helical conformation of the n+1 residue. The Y-axis in the middle panel (D2) is the free-energy barrier at the near-attack conformation defined by the ψ & χ angles. The Y-axis in the rightmost panel (D3) is the time fraction when the ASP/ASN residue remains solvent accessible. The red cut-off markers indicate the values that determine if a residue is prone to degradation or not. The blue shaded regions indicate the false negative predictions (experimentally observed to degrade but classified as non-reactive) and the grey shaded region indicates the false positives (predicted reactive but not evidenced experimentally)

To identify the relevant conformations that enable a nucleophilic attack distance, we sampled the free energy space of ψ & χ angles in Ace-GGNAG-Nme pentapeptide using Metadynamics simulations. By applying an additive historical bias, Metadynamics pushes the system to explore conformations that are kinetically limited in an equilibrium simulation. There are six low energy states in the ψ & χ space (Figure 5b), of which two regions are coincident with the near attack conformation distance (Figure 5c).

With this information, we looked at the correlation between the near attack side-chain conformation and the experimental rates reported in the adimab dataset. As with the backbone conformation, we looked at the FES from the equilibrium MD simulation. Figure 6, D2 shows that when the side-chain is not in a near-attack conformation (i.e. Free Energy > 1kcal/mol), the reaction rates largely remain below the 5% cutoff, with a few exceptions. Otherwise, the residue may or may not be reactive, as suggested by the large number of data points below and above 5% for Free Energy < 1kcal/mol. Therefore, the side chain conformation indeed proved to be a useful descriptor in identifying reactive residues (Figure 6, D2). The overall accuracy of the prediction was at 40%, but when used in combination with the backbone conformation rule, i.e. both the acidic conformation and the near attack conformations are energetically feasible, the accuracy of the classification model improves to 75% (Table 1).

#### Solvent Accessibility

Water plays an important role in the deamidation and isomerization reaction as a proton donor/acceptor. Both the base catalyzed deprotonation and hydrolysis of succinimide intermediate to ASP/iso-ASP is only feasible in the presence of water molecules (Figure 1). In previous studies, solvent accessibility weakly predicted the reaction propensity.[19, 20, 33] These previous calculations were done on static protein structures that lacked proper structural relaxations that may affect solvent accessibility calculations. Our solvent accessibility calculations utilize a dynamic approach in combination with intuitions about the reaction. Here we calculated the solvent accessibility at a residue level. In order to capture the dynamic aspect of the solvent occupancy, we represented solvent accessibility as a fraction of time in which the ASN/ASP residue remained accessible vs inaccessible (see methods). Using this value, we once again sought to classify the ASN/ASP residues in the adimab set as reactive vs non-reactive. Figure 6, D3 shows that solvent inaccessible side-chains (time fraction of solvent exposure < 75%) are indeed protected from the reaction. However, a large number of both reactive and non-reactive cases are seen in the solvent accessible region (time fraction of solvent exposure > 75%). The overall accuracy in predicting the reactivity using SASA as a single descriptor is 39%, which is no better than the performance of the near attack conformation metric.

#### Combined Model

As discussed above, deamination and isomerization reactions can be described in the three consecutive steps: deprotonation, cyclization and hydrolysis. Utilizing a multiscale computational approach, we were able to test the propensity of degradation in the adimab dataset using these physics-based metrics. Since all three steps are essential for the reaction, we sought to check if a combination of the three metrics would improve the classification accuracy: D1) deprotonation – as represented by the free-energy profile of ϕ, ψ angles of the n+1 residue recovered from the MD simulation, D2) cyclization – as represented by the free-energy profile of χ, ψ angles also recovered from the MD simulation, and D3) hydrolysis – as represented by the solvent accessibility calculated as the fraction of solvent occupied state in MD trajectory. The decision tree that we used to classify if an ASN/ASP residue is likely to degrade is shown in Figure 7. The adimab dataset discussed herein is disproportionately distributed toward true negatives (non-reactive) vs true positives (reactive), so this presented a challenge to interpreting common statistical measures such as accuracy. We instead chose to use balanced accuracy, Matthew’s correlation coefficient (MCC) and Cohen’s Kappa score (CKS) as the statistic to compare the three individual descriptors as well as combinations. We also created a naïve predictor that would predict all the residues as non-reactive; this naïve predictor will score 0 on both the MCC and CSK statistic. As can be seen from Table 1, the physics-based classifier model based on the three descriptors offers an overall balanced accuracy of 83% in predicting the isomerization and deamidation sites. All three descriptors have a better MCC and CKS score compared to the naïve predictor. Similarly, any combination of two descriptors performed moderately better than a single predictor. The combined model using all three descriptors is better at predicting non-reactive residues over reactive residues, which could be useful in a therapeutic manufacturing setting where the cost of predicting a reactive residue as non-reactive is higher. This combined mechanistic model acts as a good screening criterion in identifying antibodies that are prone to chemical degradation via deamidation or isomerization.

**Figure 7.**
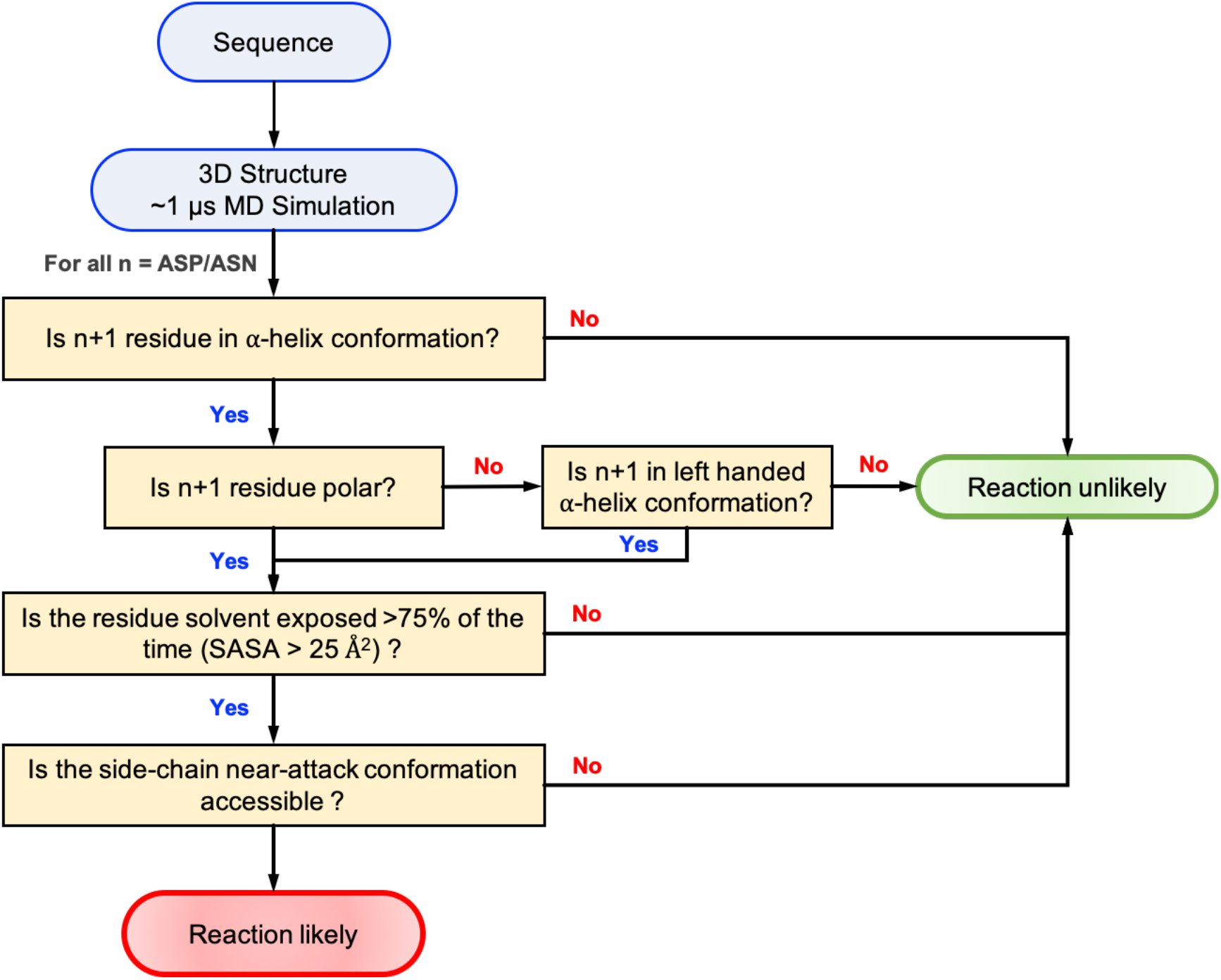
Flowchart showing the decision scheme used to determine if a given ASN/ASP residue is likely to degrade in folded proteins.

## Conclusion

Spontaneous chemical degradation at the ASN/ASP residues is an important problem that affects the developability of the protein therapeutics. Typically, the issues of chemical degradation are encountered at later stages in the development at which point iterative protein-engineering efforts are employed to mitigate these modifications. Such iterative efforts prolong development timelines and delays lifesaving therapeutics reaching patients. Multiple computational strategies ranging from QM calculation to machine learning models have been proposed to predict degradation in ASP/ASN. Very few such tools (if any) generalize for high-throughput screening. We leveraged prior theoretical and experimental findings on the ASN/ASP degradation reactions to carry out multiscale analysis to comprehensively evaluate the chemical degradation in 1039 residues (498 ASP and 541 ASN) spanning 131 therapeutic mAbs. In doing so we have made some interesting findings: 1) sidechain steric interactions of the n+1 residue affected the proton affinity of backbone amide in the right-handed α-helix conformation, 2) NG and DG sites as well as the reactive N/DX sites are preferentially observed in the left-handed α-helix conformation that correlates with low proton affinity, and 3) there are specific regions of the sidechain conformational space are both energetically favorable and in a near-attack conformation. Utilizing an experimentally validated truth set we have been able to connect molecular intuition from the reaction mechanism to a real-life scenario. Utilizing just three descriptors that are fundamental to the protein structure one can heuristically classify the likelihood of the protein to undergo chemical degradation at the ASN/ASP sites. Although we have exclusively looked at mAbs in this study, we believe that the structural and chemical insights described herein are broadly applicable and generalizable to all proteins and protein-based therapeutics.

## Methods

### Experimental dataset

The validation data contained 498 ASP and 541 ASN residues in the CDR loops of 131 therapeutic mAbs published by Lu et al.[34] The isomerization events were analyzed at low pH stress (pH 5.5) for 2 weeks at 40 °C. A total of 31 isomerization sites were identified with total modification measurements ranging from to 44.0%. The deamidation data was obtained at high pH stress (pH 8.5) for 1 week at 40 °C. A total of 39 deamidation sites were reported with degradation levels ranging from 2.0% to 67.7%. We used a threshold cutoff of 5.0% for both isomerization and deamidation events to classify reactive versus non-reactive.

### QM calculations

Semi-empirical QM (SQM) calculations to calculate proton affinity were performed using GAUSSIAN09[50] and MOPAC2016[51] software packages. To rule out any uncertainties in the proton affinity calculations, we carried out DFT calculation of proton affinity for N-formyl-glycinamide at M062X/6-311++G(d,p) level of theory in both gas phase and aqueous phase continuum SMD solvent environment.[52] The gas phase calculations where then repeated in MOPAC with PM6-D3H4,[53] PM7 and RM1 semi-empirical methods (SI).[52] The regions of proton affinity qualitatively matched across DFT and SQM levels of theory albeit with variations in the energy values as expected due to variations in the theory and the Hamiltonian. Throughout the manuscript we tried to make only qualitative inferences from the QM calculations and we refrained from interpreting the exact energy as that is beyond the scope of the current study.

Dipeptides of the Ace-NX-Nme and Ace-DX-Nme as well as their corresponding deprotonated anions (where X can be single letter code of all 20 amino acids) were generated using openbable.[54] The generated structures were subject to geometry optimization under PM6 level of theory to arrive at an initial structure. A 2D PES scan at PM6-D3H4, PM7 and RM1 levels of theory. The scan was performed at 10° intervals spanning the entire Ramachandran region. A total of 1369 data points were scanned for each dipeptide and its corresponding anion. At each step geometry optimization was requested to adhere to TIGHT convergence criterion while the angles being scanned would be kept constant. The resulting PES were used to arrive at the proton affinity expressed in the heat of formation for the forward reaction as described in Figure 2. The discrete energy points were interpolated and plotted using linear spline interpolation as implemented in the matplotlib[55] python package.

### PDB Library

We downloaded the pdb file repository of all the antibody (and antibody like Fab, scFV, ect…) that have crystal structure deposited in the PDB from SAbDab (The Structural Antibody Database http://opig.stats.ox.ac.uk/webapps/newsabdab/sabdab/).[56, 57] We arrived at 3493 structures which were then cleaned, energy minimized and isolated from any other ligands. Using these structures, we calculated backbone dihedral angles of all Nx and Dx residues from the Fab domain. All the PDB handling was performed using MDanalysis[58] and PyTraj[59] python tools. The backbone dihedral angle distribution from this data was used for calculating 2D kernel density estimation using the SciKit-Learn python package.[60]

### KDE calculation

In order to compare the KDEs that were calculated from the PDB database and the backbone dihedral angles observed in the MD simulations, we first classified each N/Dx site as reactive or non-reactive based on the deprotonation metric. We collected the ϕ & ψ angle distribution for all these sites across all MD trajectories. This data was used to calculate the second kernel density estimation plot in Figure 4, using the same SciKit-Learn package as before.

### Molecular Dynamics Simulation

Molecular Operating Environment (MOE)[61] software from Chemical Computing Group was used to homology model each of the fragment antigen-binding (Fab) domain of all the antibodies accessed in the adimab set. The Fab structures were energy minimized to remove any steric clashes. The energy minimization was performed using SANDER program in Amber2015.[62] The protonation states at pH 5.5 and 8.5 were assigned using the PDB2PQR tool. The two pH values mimic the buffer conditions in which the reactivity was estimated in the adimab experiment.

Energy minimized Fab structures were then prepared for equilibrium MD simulations by solvating the structure in a cubic box of TIP3P explicit water (model).[63] It was ensured such that the Fab was positioned at a minimum 10Å distance from the edge of the box. The system was then adjusted to have a net neutral charge by adding Na+ and Cl-counter ions. Hydrogen Mass Repartitioning was performed on the solute atoms to enable long simulation time steps of 4fs.[64]

The GPU implementation of Amber 2015 MD software package with the SPFP precision model was used for the MD simulation. The simulations proceeded by relaxing the system with 2000 steps of conjugate-gradient energy minimization. Harmonic restraining potentials with the force constant of 10 kcal/mol/Å^2 were imposed to restrain the solute to its initial structure. Post which the system was brought to an NPT ensemble with the pressure maintained at 1 atm and the thermostat set to 300K over the course of 200 ps, while the protein was restrained using a harmonic positional restraint of 10 kcal/mol/Å^2. The system was then equilibrated for 1 ns with a restraint force constant of 1 kcal/mol/Å^2 on the protein structure maintaining the pressure and temperature at the same levels. The system was then subject to equilibrium MD simulation at NPT ensemble without any restraints at a time step of 4 fs. Long range interactions were cut off at 9Å and restricted range limited interactions. Coulombic interaction was calculated using the Particle Mesh Ewald algorithm for long-range electrostatics. The thermostat was maintained at 300K with Langevin dynamics and collision frequency set to 1 ps^−1^. The simulation was carried out for 500 ns during which the SHAKE algorithm was applied to constrain all bonds involving hydrogen atoms. Two other replicates of the 500 ns simulation were performed instantiated with a different random number to bring the combined sampling time to 1.5 *μs*. Trajectory snapshots were saved at 10 ps intervals and used for downstream analysis. The combined trajectories from three simulations were used for the analysis.

CPPTRAJ software in AmberTools was used to analyze the trajectories. The free energy surface (FES) in the space of dihedral angles were calculated from bin populations using Gi=-kB T ln(Ni/Nmax), where kB is Boltzmann’s constant, T is the temperature, Ni is the population of bin i and Nmax is the population of the most populated bin. Bins with no population were given an artificial barrier equivalent to a population of 0.5. Solvent accessible surface area (SASA) was calculated using the MSMS package[65] using a probe radius of 1.4 and triangle density of 3.

### Advanced Sampling Using Metadynamics

Metadynamics simulations were carried out using the GROMACS2018[66] MD engine patched to interface with PLUMED.[67] Metadynamics was performed in the well-tempered[68] scheme along the conformational space of ψ & χ angles of the pentapeptide Ace-GGNAG-Nme. The simulations utilized the same forcefield and MD parameters with 2 fs time steps. Bias was added at every 500 steps with a sigma of 0.1 and bias of 1.0 kcal/mol. The simulation was carried out for 1 μs and the trajectory was analyzed after unbiasing the simulation using the standard reweighting[69] procedure implemented in PLUMED. The free energy surface and corresponding distance distribution was plotted using matplotlib.

## Acknowledgements

The authors would like to thank Robert Kelly, Vikas Sharma and Sreedhara Alavattam for their engaging discussions. We would like to thank Benjamin Sellers for his valuable comments and suggestions on the manuscript.

## Supporting Information

The supporting information provides the proton affinity calculations and the corresponding KDE plots for other dipeptides.

## Contact

All correspondence and data requests should be directed to the corresponding author.

