## Supplemental Information for "Divining Deamidation and Isomerization in Therapeutic Proteins: Effect of Neighboring Residue"

### 2-acetamido-N-methylacetamide

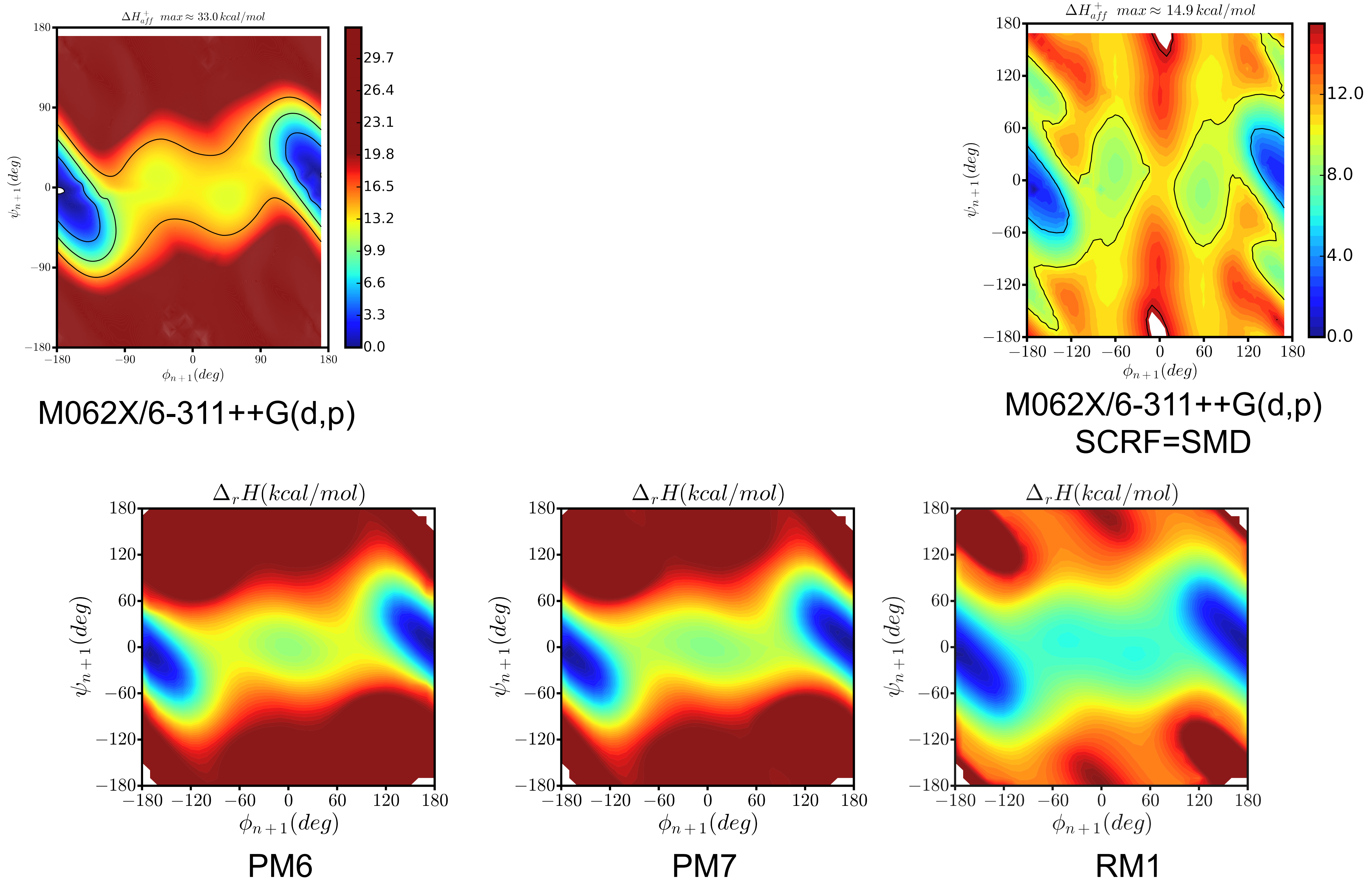

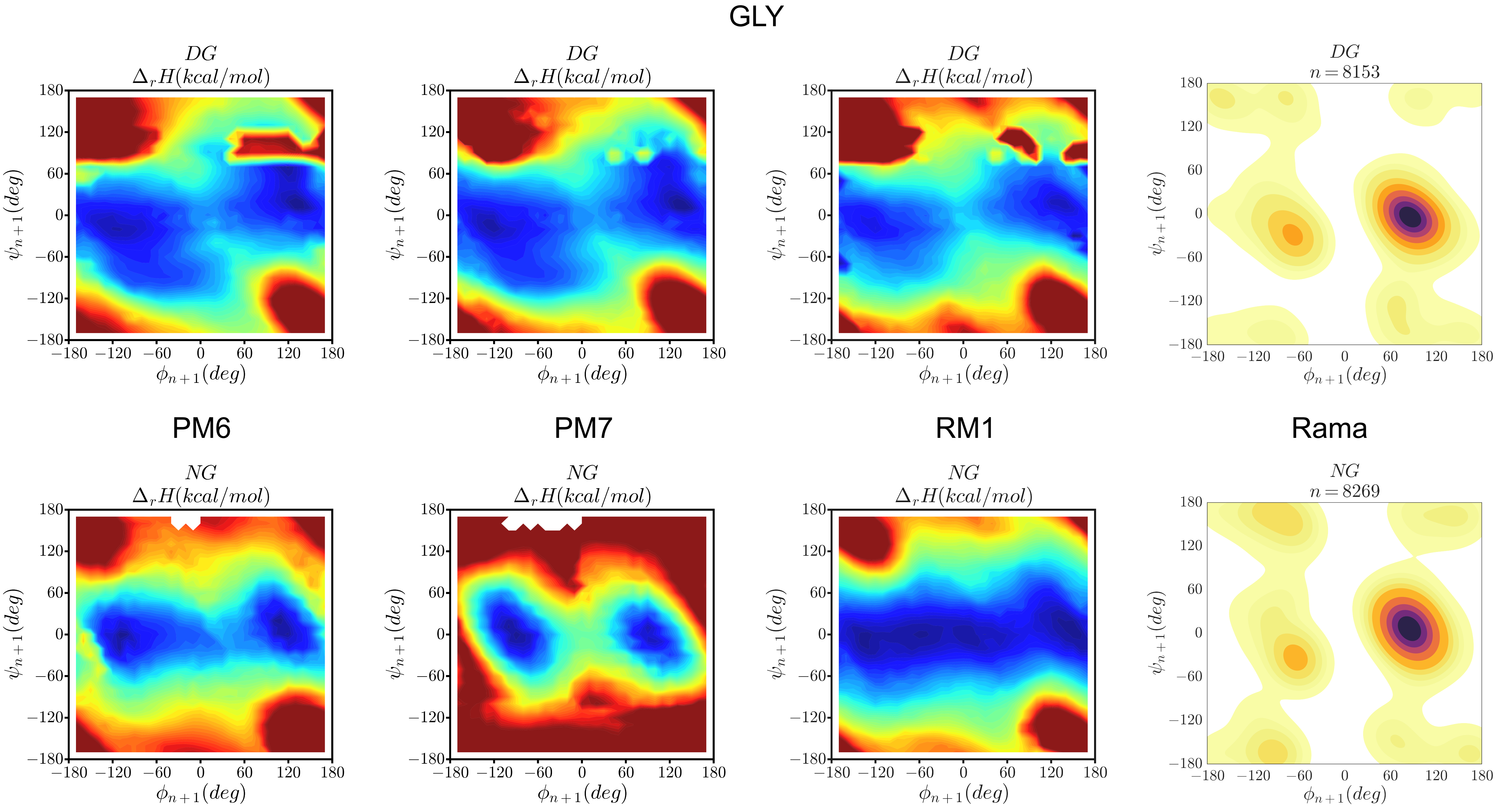
GLY
DG
$\Delta_r H(kcal/mol)$
DG
$\Delta_r H(kcal/mol)$
DG
$\Delta_r H(kcal/mol)$
DG
$n = 8153$
PM6
PM7
RM1
Rama
NG
$\Delta_r H(kcal/mol)$
NG
$\Delta_r H(kcal/mol)$
NG
$\Delta_r H(kcal/mol)$
NG
$n = 8269$
$\psi_{n+1}(deg)$
$\phi_{n+1}(deg)$
−180 −120 −60 0 60 120 180

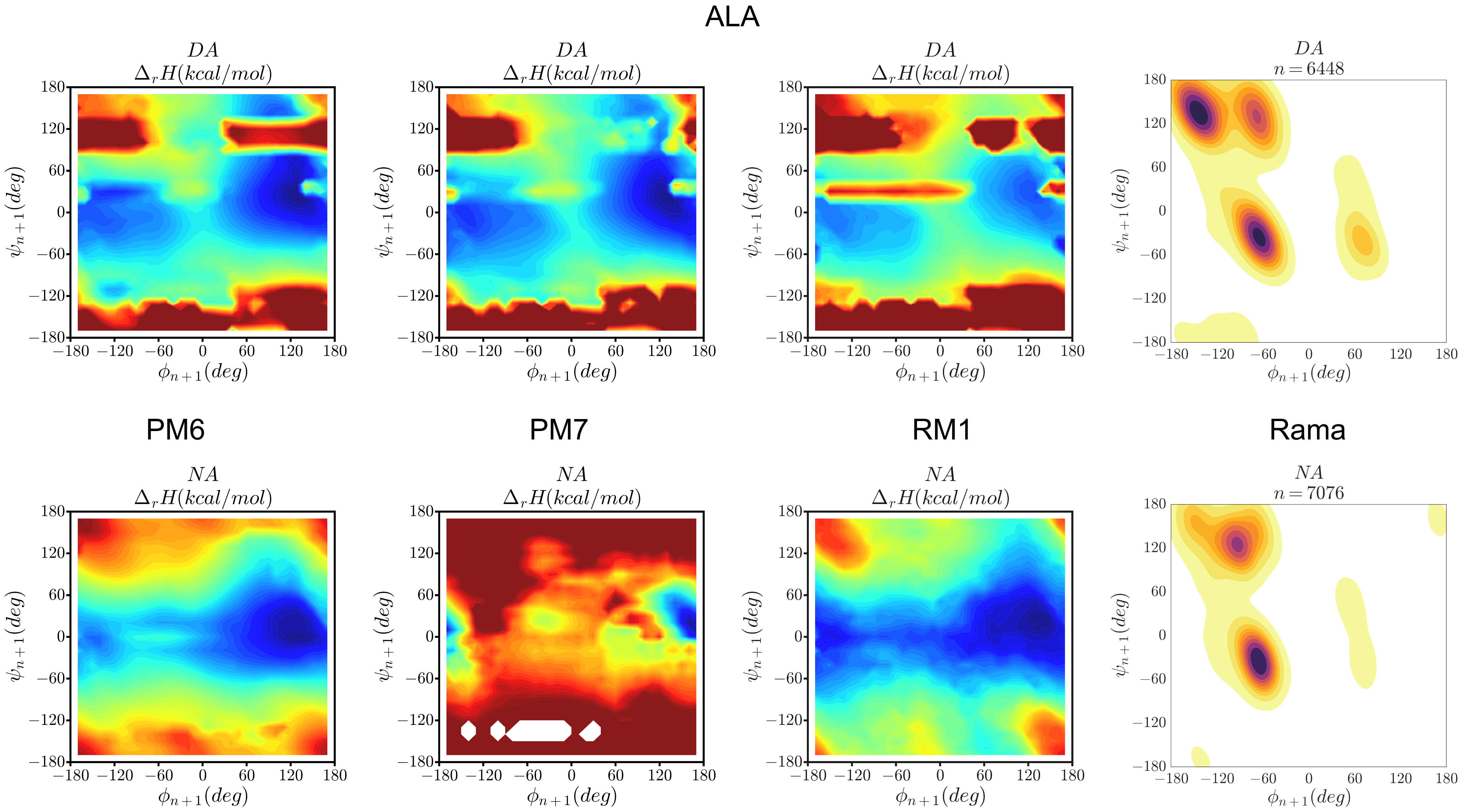

ALA
PM6
PM7
RM1
Rama
DA
NA
$\Delta_r H(kcal/mol)$
$n = 6448$
$n = 7076$
$\phi_{n+1}(deg)$
$\psi_{n+1}(deg)$
−180
−120
−60
0
60
120
180

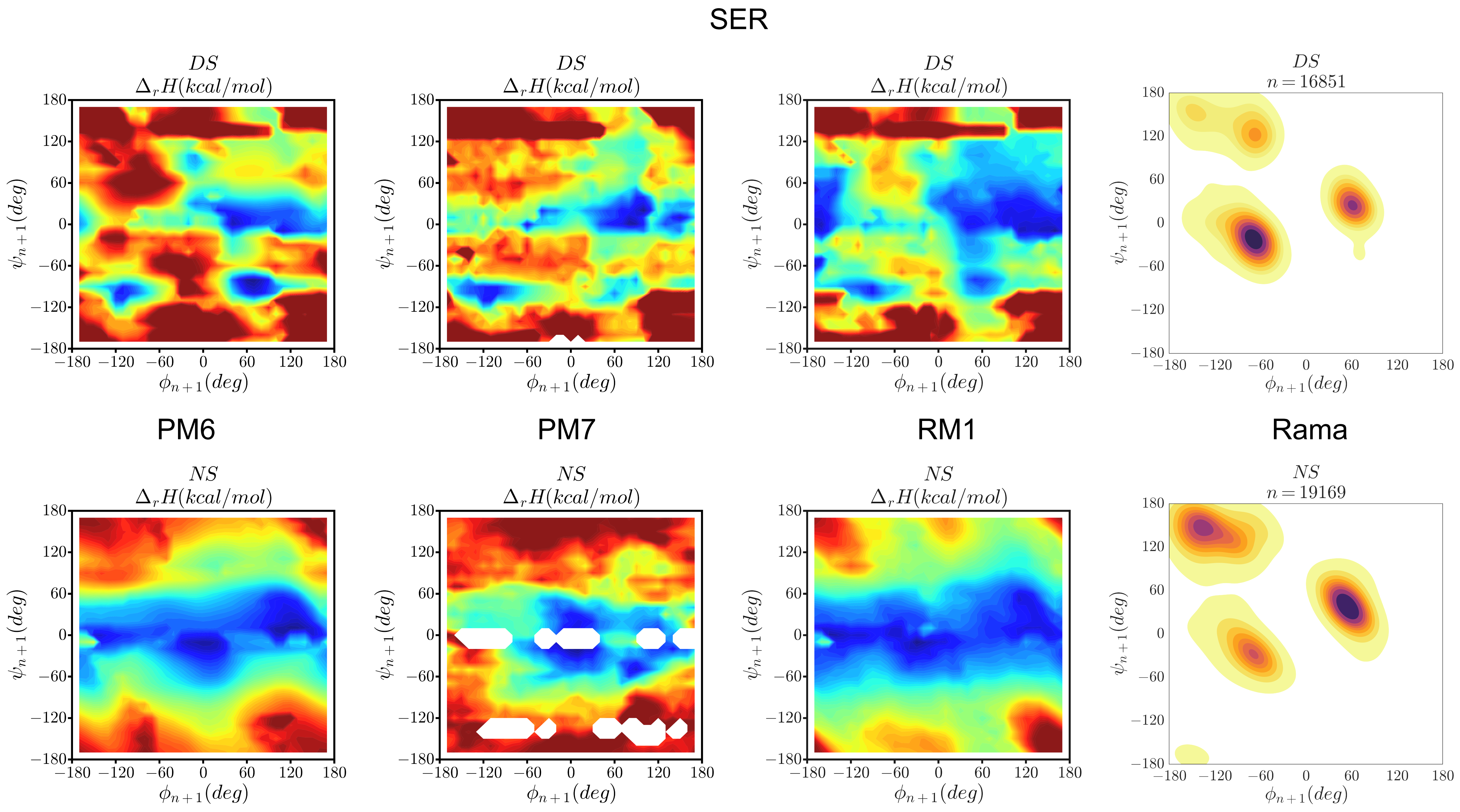

SER
DS
$\Delta_r H (kcal/mol)$
DS
$n = 16851$
PM6
PM7
RM1
Rama
NS
$\Delta_r H (kcal/mol)$
NS
$n = 19169$
$\psi_{n+1} (deg)$
$\phi_{n+1} (deg)$
180
120
60
0
−60
−120
−180

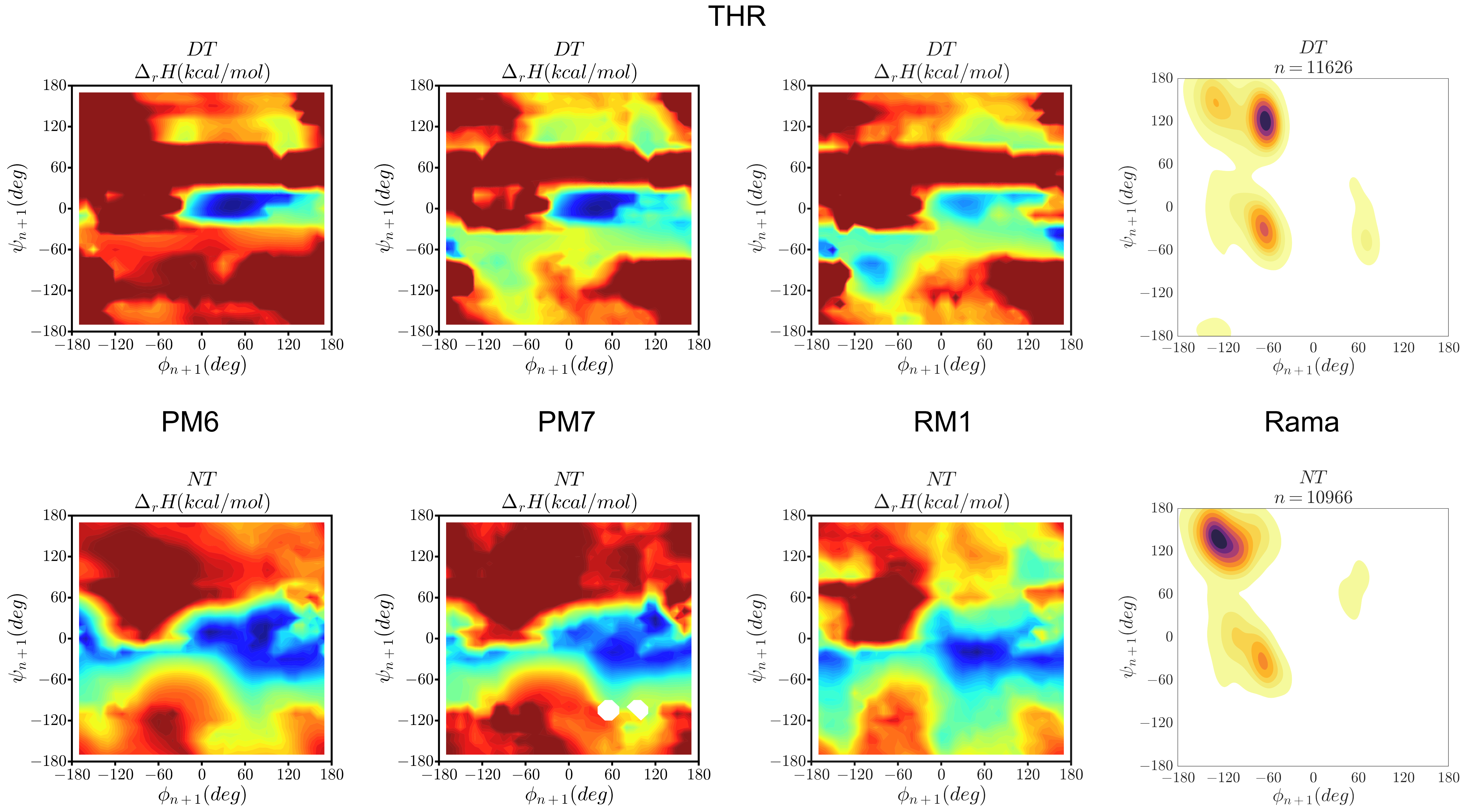

THR
DT
$\Delta_r H(kcal/mol)$
NT
$n = 11626$
$n = 10966$
PM6
PM7
RM1
Rama
$\phi_{n+1}(deg)$
$\psi_{n+1}(deg)$

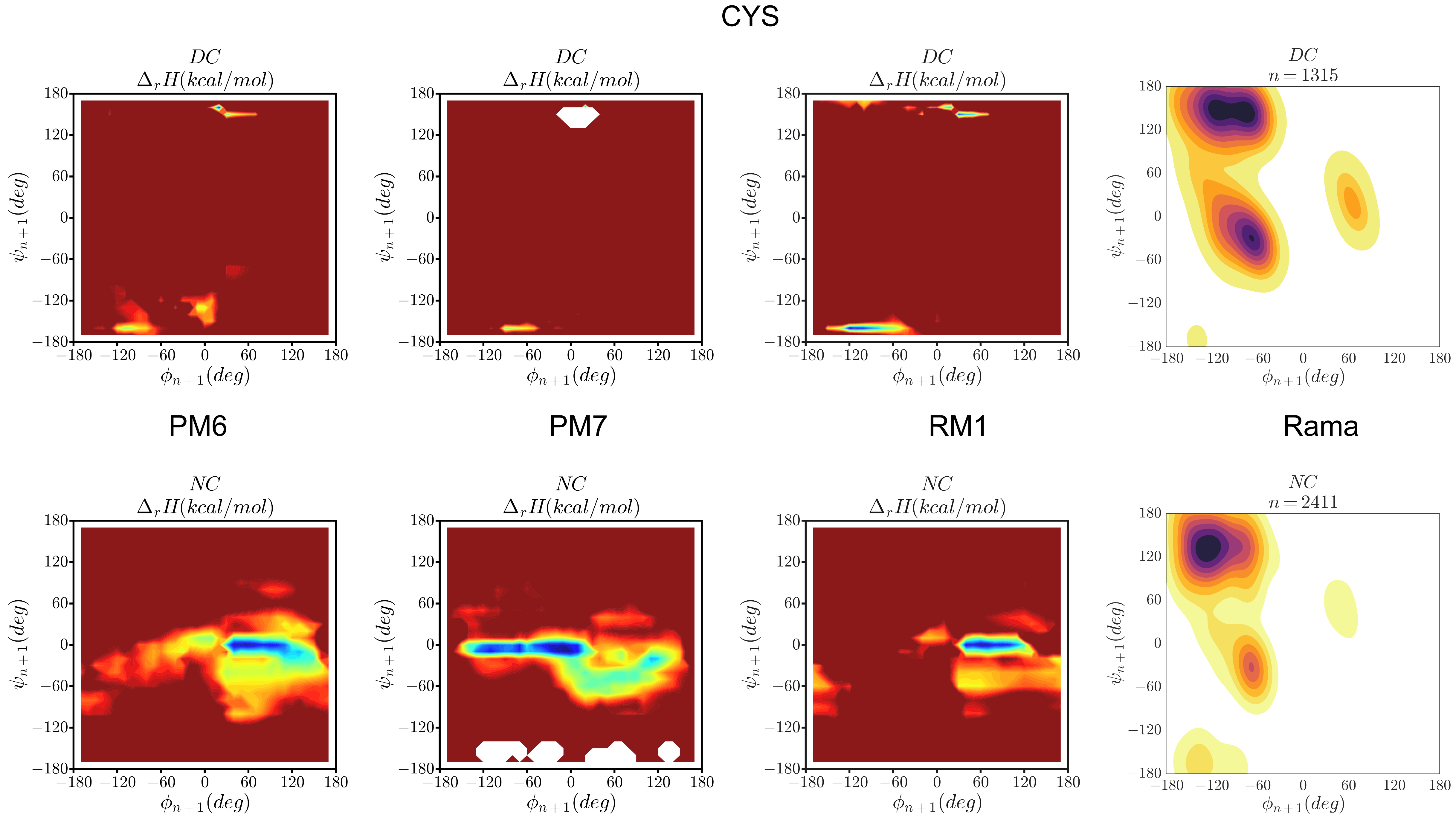
CYS
DC
$\Delta_r H(kcal/mol)$
DC
$n = 1315$
PM6
PM7
RM1
Rama
NC
$\Delta_r H(kcal/mol)$
NC
$n = 2411$
$\psi_{n+1}(deg)$
$\phi_{n+1}(deg)$
−180
−120
−60
0
60
120
180

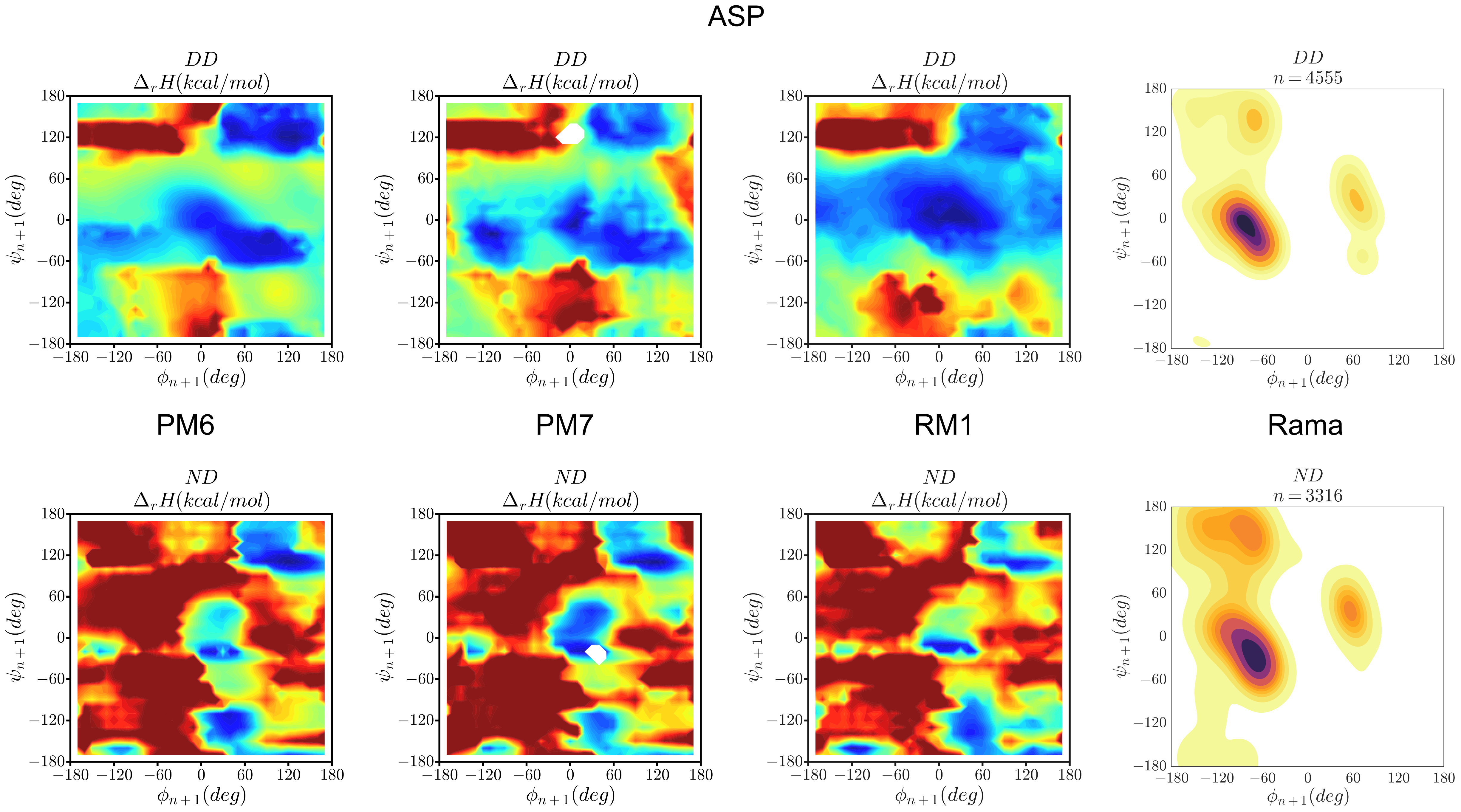
ASP
DD
ΔrH(kcal/mol)
n = 4555
PM6
PM7
RM1
Rama
ND
ΔrH(kcal/mol)
n = 3316
ψn+1(deg)
φn+1(deg)
−180 −120 −60 0 60 120 180

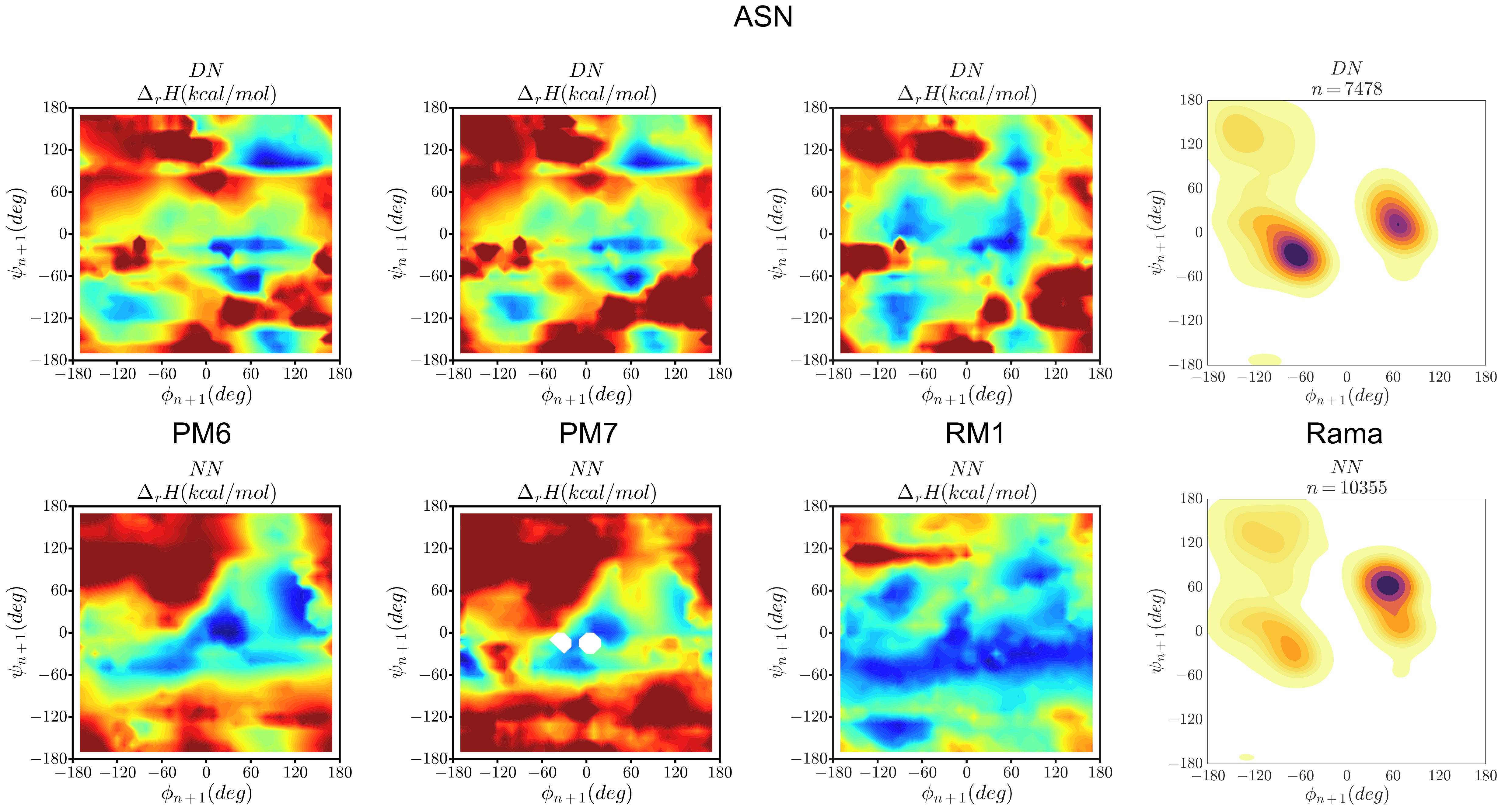

ASN
DN
$\Delta_r H(kcal/mol)$
NN
$\Delta_r H(kcal/mol)$
DN
$n = 7478$
NN
$n = 10355$
PM6
PM7
RM1
Rama
$\phi_{n+1}(deg)$
$\psi_{n+1}(deg)$
−180
−120
−60
0
60
120
180

### GLN

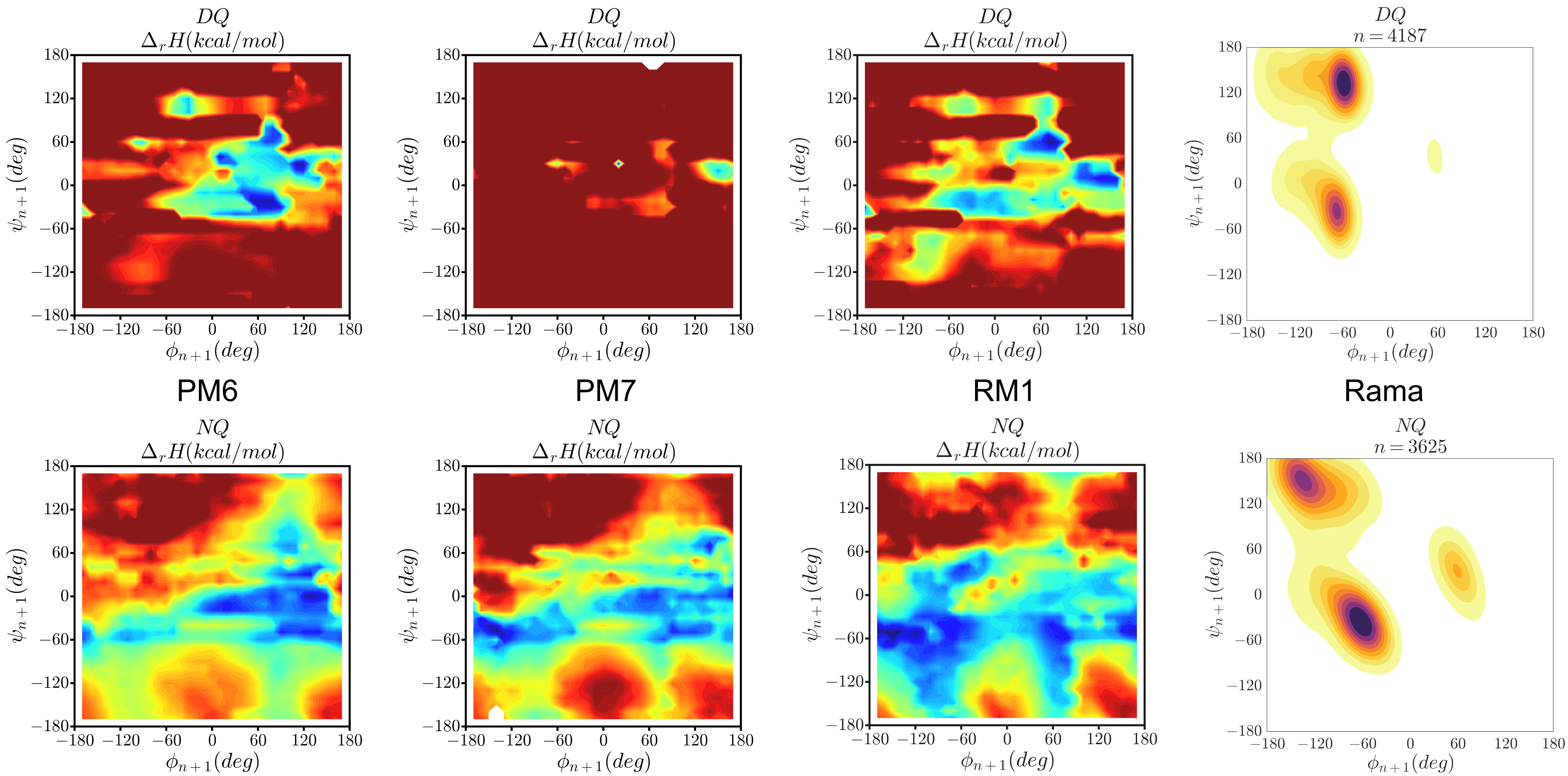

GLN
DQ
$\Delta_r H(kcal/mol)$
NQ
$n = 4187$
$n = 3625$
PM6
PM7
RM1
Rama
$\phi_{n+1}(deg)$
$\psi_{n+1}(deg)$
−180
−120
−60
0
60
120
180

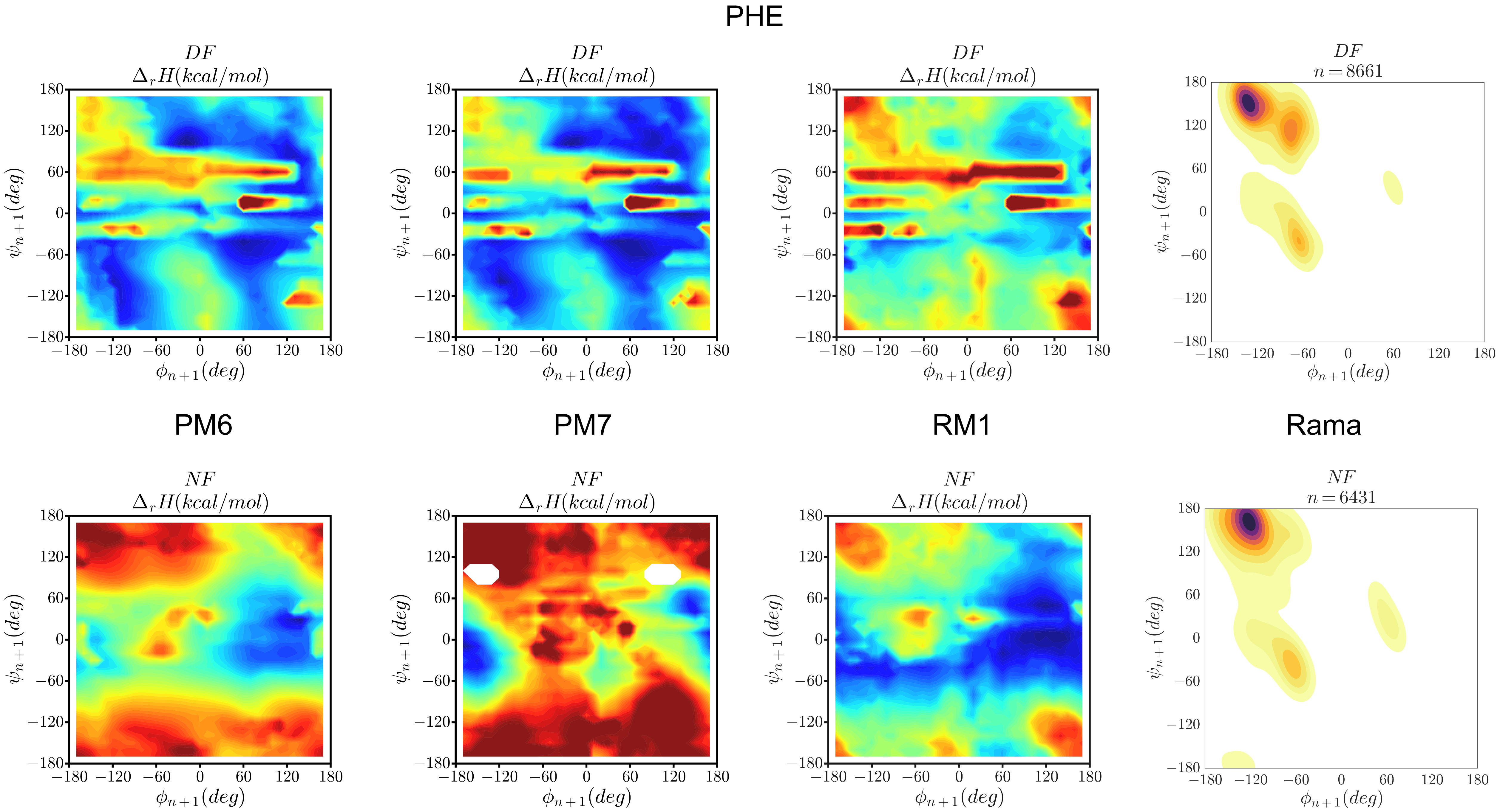
PHE
PM6
PM7
RM1
Rama
DF
NF
$\Delta_r H (kcal/mol)$
$n = 8661$
$n = 6431$
$\phi_{n+1}(deg)$
$\psi_{n+1}(deg)$
−180
−120
−60
0
60
120
180

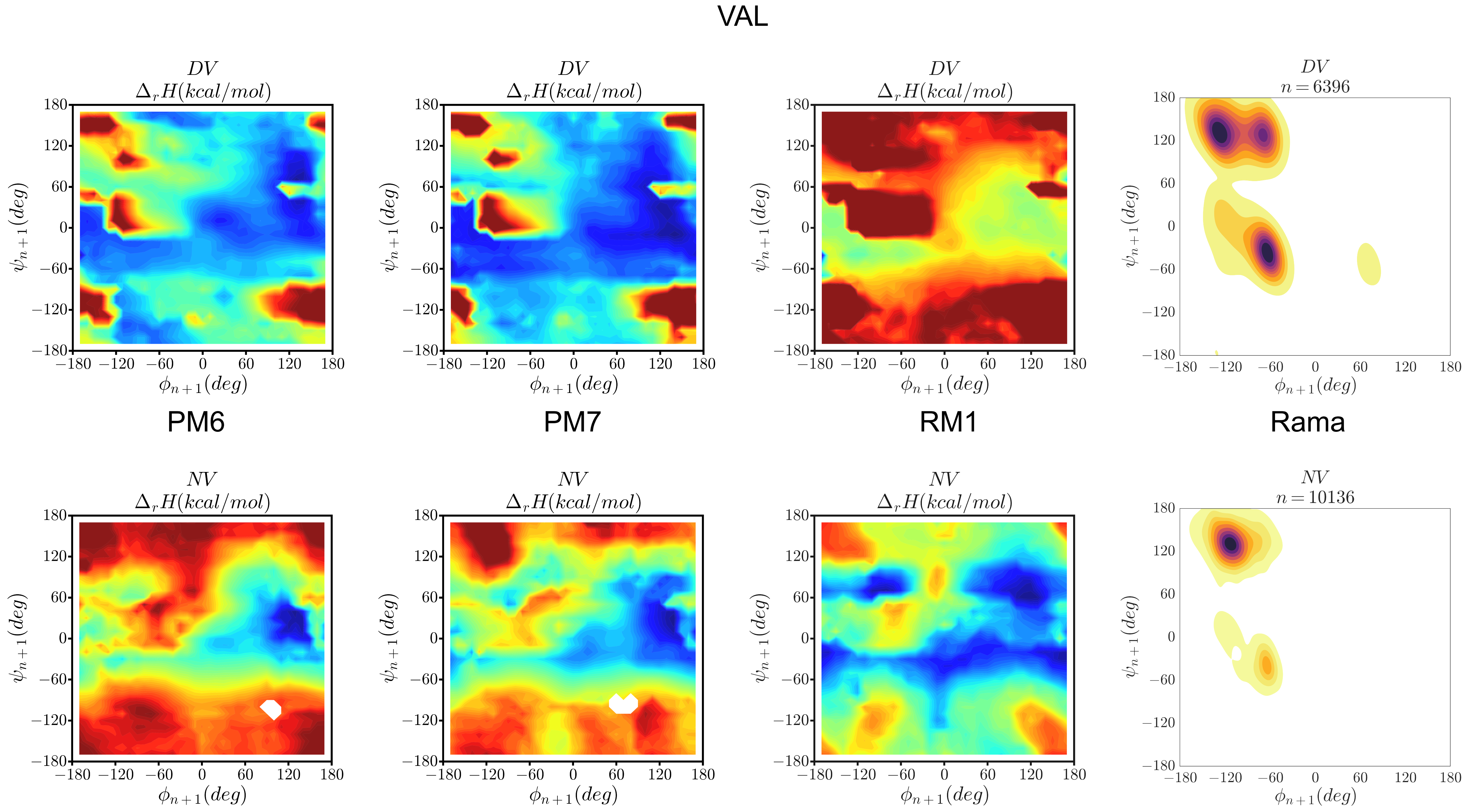
VAL
DV
$\Delta_r H(kcal/mol)$
$\psi_{n+1}(deg)$
$\phi_{n+1}(deg)$
PM6
PM7
RM1
Rama
$n = 6396$
NV
$n = 10136$
180
120
60
0
−60
−120
−180

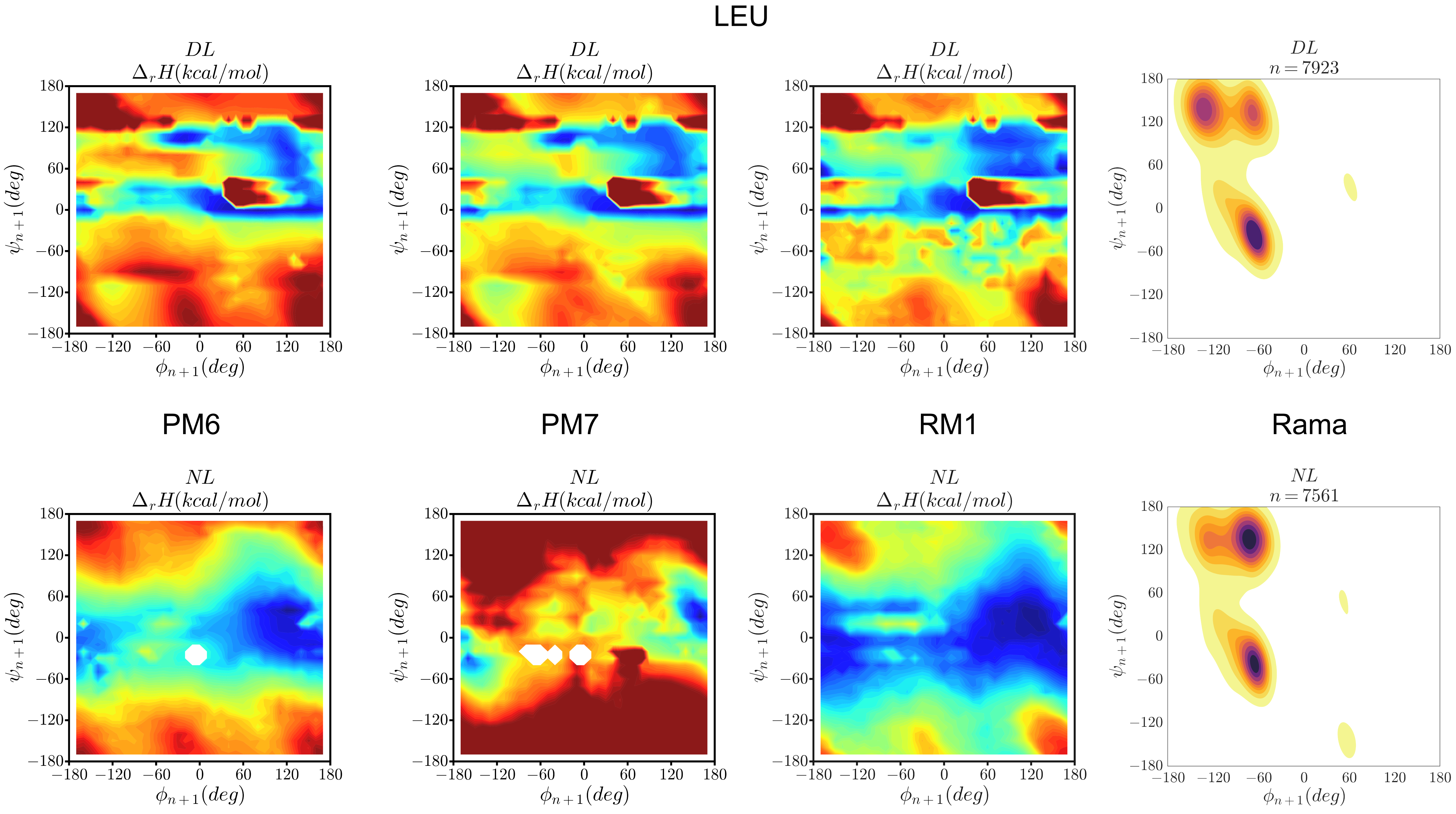

LEU
PM6
PM7
RM1
Rama
DL
Δ_rH(kcal/mol)
n = 7923
NL
n = 7561
ψ_{n+1}(deg)
φ_{n+1}(deg)
180
120
60
0
−60
−120
−180

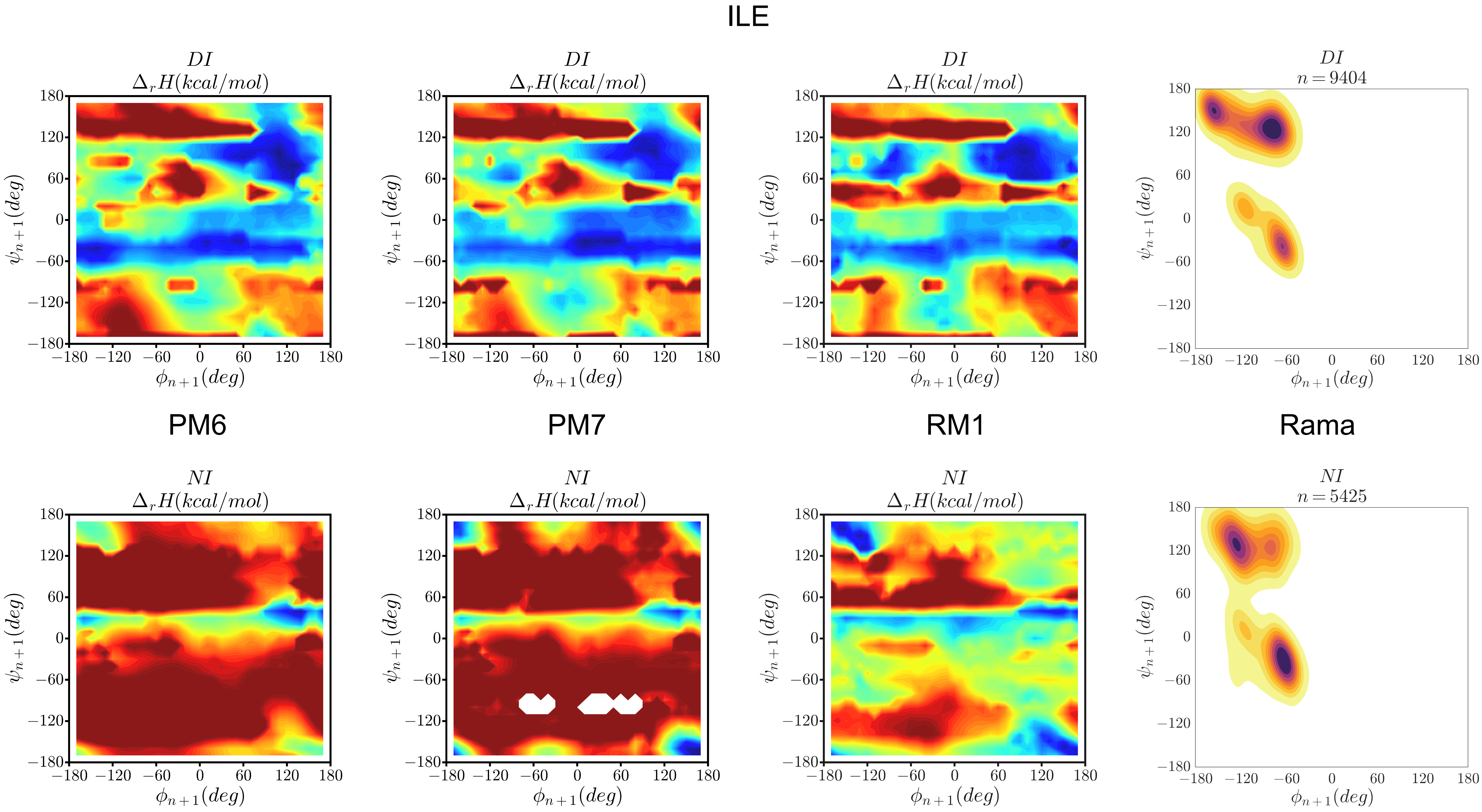
ILE
DI
$\Delta_r H(kcal/mol)$
PM6
PM7
RM1
Rama
DI
$n = 9404$
NI
$\Delta_r H(kcal/mol)$
NI
$n = 5425$
$\psi_{n+1}(deg)$
$\phi_{n+1}(deg)$
180
120
60
0
−60
−120
−180

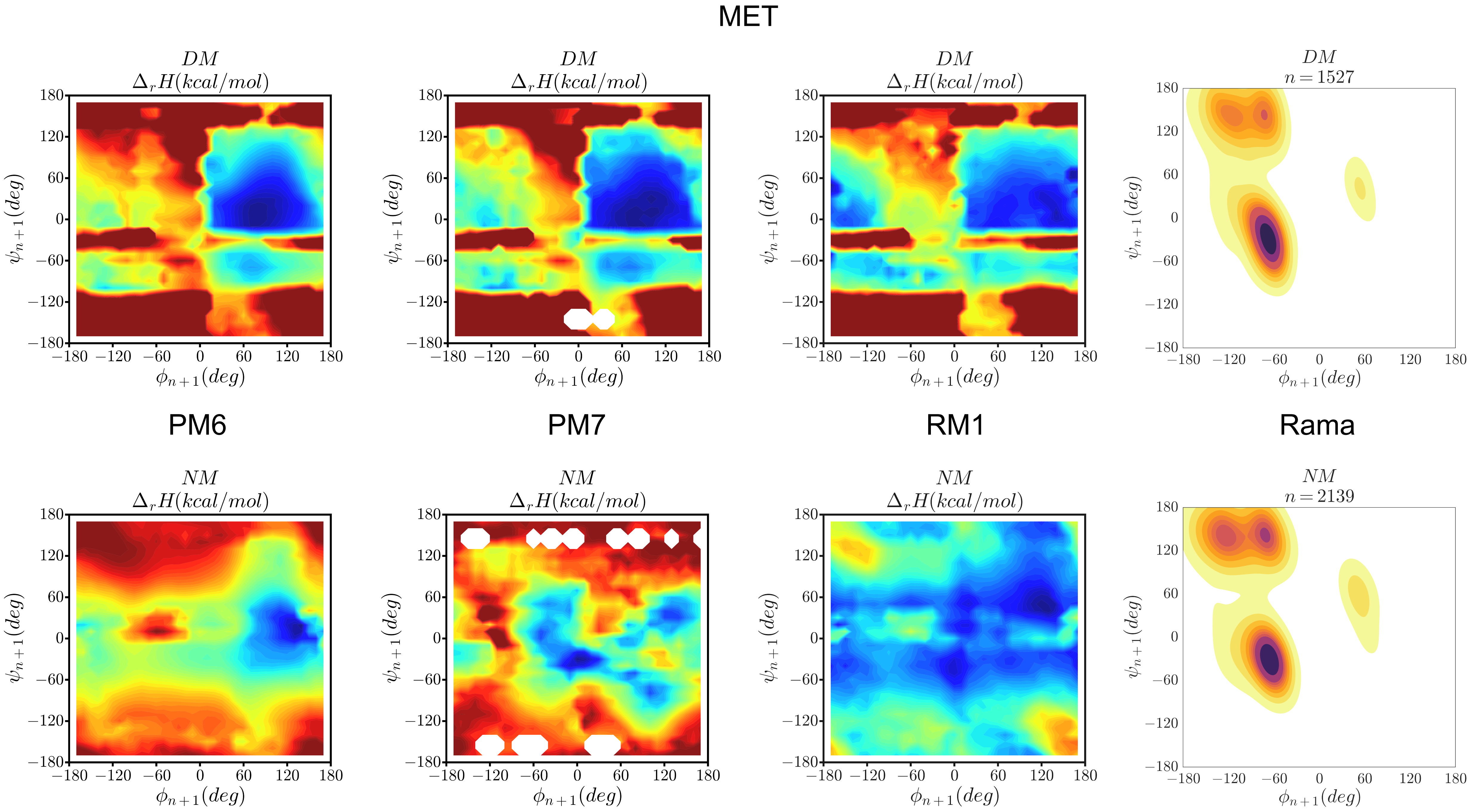

MET
DM
$\Delta_r H(kcal/mol)$
DM
$n = 1527$
PM6
PM7
RM1
Rama
NM
$\Delta_r H(kcal/mol)$
NM
$n = 2139$
$\psi_{n+1}(deg)$
$\phi_{n+1}(deg)$
